# Human skin rejuvenation via mRNA

**DOI:** 10.1101/2024.11.12.623261

**Authors:** Li Li, Zhengkuan Tang, Xavier Portillo, Rohit Arora, Madeleine Yang, Esther Mintzer, Zhenrui Zhang, Stefano Sol, Fabiana Boncimino, Patrick Han, Michelle Dion, Emery Di Cicco, Dorothy D. Majewski, Asaf Ashkenazy Titelman, Alexandru M. Plesa, Michael Shadpour, Shannon M. McNamee, Paula Restrepo, Andrew L. Ji, Juseong Lee, Yue Zheng, Xiaolong Li, Yu Wang, Jenny M. Tam, Jean S. McGee, Yu Shrike Zhang, Natalie Artzi, George F. Murphy, Anna Mandinova, Thomas S Kupper, Sai Ma, George M. Church

**Affiliations:** Department of Genetics, Harvard Medical School, Boston, MA, USA; Wyss Institute for Biologically Inspired Engineering, Harvard University, Boston, MA, USA; Division of Engineering in Medicine, Department of Medicine, Brigham and Women’s Hospital, Harvard Medical School, Boston, MA, USA; Ragon Institute of MGH, MIT and Harvard, Cambridge, MA, USA; Cutaneous Biology Research Center, Massachusetts General Hospital and Harvard Medical School, Charlestown, MA, USA; Harvard-MIT Division of Health Sciences and Technology, Massachusetts Institute of Technology, Cambridge, MA, USA; Institute for Medical Engineering and Science, Massachusetts Institute of Technology, Cambridge, MA, USA; Department of Biological Engineering, MIT, Cambridge, MA, USA; Harvard Stem Cell Institute, Cambridge, MA, USA; Department of Pathology, Brigham and Women’s Hospital, Harvard Medical School, Boston, MA, USA; Graduate School of Biomedical Sciences, Icahn School of Medicine at Mount Sinai, New York, NY, USA; Black Family Stem Cell Institute, Icahn School of Medicine at Mount Sinai, New York, NY, USA; Kimberly and Eric J. Waldman Department of Dermatology, Icahn School of Medicine at Mount Sinai, New York, NY, USA; Tisch Cancer Institute, Icahn School of Medicine at Mount Sinai, New York, NY, USA; Department of Oncological Sciences, Icahn School of Medicine at Mount Sinai, New York, NY, USA; Marc and Jennifer Lipschultz Precision Immunology Institute, Icahn School of Medicine at Mount Sinai, New York, NY, USA; Department of Cell, Developmental and Regenerative Biology, Icahn School of Medicine at Mount Sinai, New York, NY, USA; Nanfang Hospital, Southern Medical University, Guangzhou, China; State Key Laboratory of Immune Response and Immunotherapy & Department of Hematology, The First Affiliated Hospital of USTC, Hefei 230001, Anhui, China; MOE Key Laboratory for Cellular Dynamics, School of Life Sciences, Division of Life Sciences and Medicine, University of Science and Technology of China, Hefei 230001, Anhui, China; Institute of Digital Medicine, City University of Hong Kong, Hong Kong; Department of Dermatology, Brigham and Women’s Hospital, Boston, MA, USA; Broad Institute of Harvard and MIT, Cambridge, MA, USA; Harvard Medical School, Boston, MA, USA; Harvard Skin Disease Research Center, Harvard Medical School, Boston, MA, USA; Department of Genetics and Genomic Sciences, Icahn School of Medicine at Mount Sinai, New York, NY, USA

**Keywords:** skin rejuvenation, mRNA treatment, skin stem cell, aging

## Abstract

Aging is characterized by a gradual decline in function, partly due to accumulated molecular damage. Human skin undergoes both chronological aging and environmental degradation, particularly UV-induced photoaging. Detrimental structural and physiological changes caused by aging include epidermal thinning due to stem cell depletion and dermal atrophy associated with decreased collagen production. Here, we present a comprehensive single-cell atlas of skin aging, analyzing samples from young, middle-aged, and elderly individuals, including both sun-exposed and sun-protected areas. This atlas reveals age-related changes in cellular composition and function across various skin cell types, including epidermal stem cells, fibroblasts, hair follicles, and endothelial cells. Using our atlas, we have uncovered basal stem cells as a highly variable population across aging, more so than other skin cell populations, such as fibroblasts. In basal stem cells, we identified ATF3 as a novel regulator of skin aging. ATF3 is a transcriptional factor for genes involved in the aging process, with its expression reduced by 20% during aging. Based on this discovery, we developed an innovative mRNA-based treatment to mitigate the effects of skin aging. After treatment with ATF3 mRNA, cell senescence decreased 25%, and we observed an over 20% increase in proliferation in treated basal stem cells. Importantly, we also found communication between keratinocytes and fibroblasts as a critical component of therapeutic interventions, with ATF3 mRNA rescue of basal cells significantly enhancing fibroblast collagen production by approximately 200%. Furthermore, we validated the efficacy of ATF3 mRNA treatment in *ex vivo* human skin and *in vivo* mouse models. In *ex vivo* human skin, microneedle-mediated delivery of ATF3 mRNA induced robust rejuvenation, expanding the basal stem-cell population and enhancing dermal ECM reconstruction. In a wound healing mouse model, ATF3 mRNA reduced scarring, demonstrating strong regenerative potential for reversing age-related skin decline. We conclude that ATF3 mRNA treatment effectively reverses the effects of skin aging by modulating specific cellular mechanisms, offering a novel, targeted approach to human skin rejuvenation.

## INTRODUCTION

Aging is a fundamental biological process marked by a progressive decline in cellular functions and driven by the accumulation of molecular damage. Skin aging manifests through both chronological and environmental processes, particularly UV-induced photoaging, leading to significant and detrimental structural and physiological changes ^1^. The complexity of these changes and the critical importance of skin as a barrier to a potentially hostile external environment highlight the urgent need for effective strategies to combat skin aging and mitigate its detrimental effects.

Various treatments have been proposed to address skin aging and related conditions, including retinoic acid application, intense pulsed light therapy, laser treatments, and biological approaches involving platelet-rich plasma and mesenchymal stem cells ^2–10^. Despite these modalities, effective treatments for skin rejuvenation remain to be elucidated, particularly ones that address both intrinsic and extrinsic mechanisms of skin aging. This gap likely relates to an incomplete understanding of the molecular changes occurring within different skin cell types during aging. The majority of previous studies have focused exclusively on aging and rejuvenation mechanisms targeting fibroblasts. However, approaches to skin aging would benefit from consideration of other skin cell populations, such as basal stem cells, affected by the aging process. Thus, the complexity of skin aging and the limitations of current treatments motivate more detailed experimental inquiry into the fundamental pathways that mediate the skin aging process.

In this study, we employed single-cell omics techniques to develop a comprehensive Human Skin Atlas, providing a detailed map of differential gene expression patterns across various ages and skin conditions. This approach enabled us to identify ATF3 in basal stem cells as a crucial regulator of skin aging. We learned that ATF3 expression decreases significantly with age, contributing to increased cell senescence and altered keratinocyte-mediated fibroblast collagen production. Building on this discovery, we developed and experimentally validated an innovative mRNA-based therapeutic approach targeting ATF3 with *in vitro* cell culture, *ex vivo* human skin, and *in vivo* murine models. This novel treatment demonstrates significant potential in reversing skin aging, addressing both intrinsic and extrinsic aging mechanisms.

## RESULTS

### A comprehensive single-cell dataset of aging human skin

To gain detailed insights into the molecular changes associated with skin aging at the single-cell level for treatment development, we conducted a comprehensive and in-depth analysis of healthy human adult skin samples across various age groups using single-cell RNA sequencing (scRNA-seq) (**Fig. 1a and Supplementary Table 1**). In total, we analyzed 16 human skin samples, representing one of the largest single-cell transcriptomic datasets currently available for human skin ^11^. The 10 mm biopsies were collected from young (Y), middle-aged (M), and old (O) donors. Since skin aging involves both intrinsic (chronological) and extrinsic (photo) aging processes, paired samples were obtained from the arm (subject to both chronological and photoaging) and the back (subject primarily to chronological aging) ^12^. Importantly, these paired biopsies were collected from two anatomically distinct sites of the same individuals, thereby minimizing inter-individual variability and providing a valuable resource for dissecting both photoaging and intrinsic aging mechanisms.

**Fig. 1.**
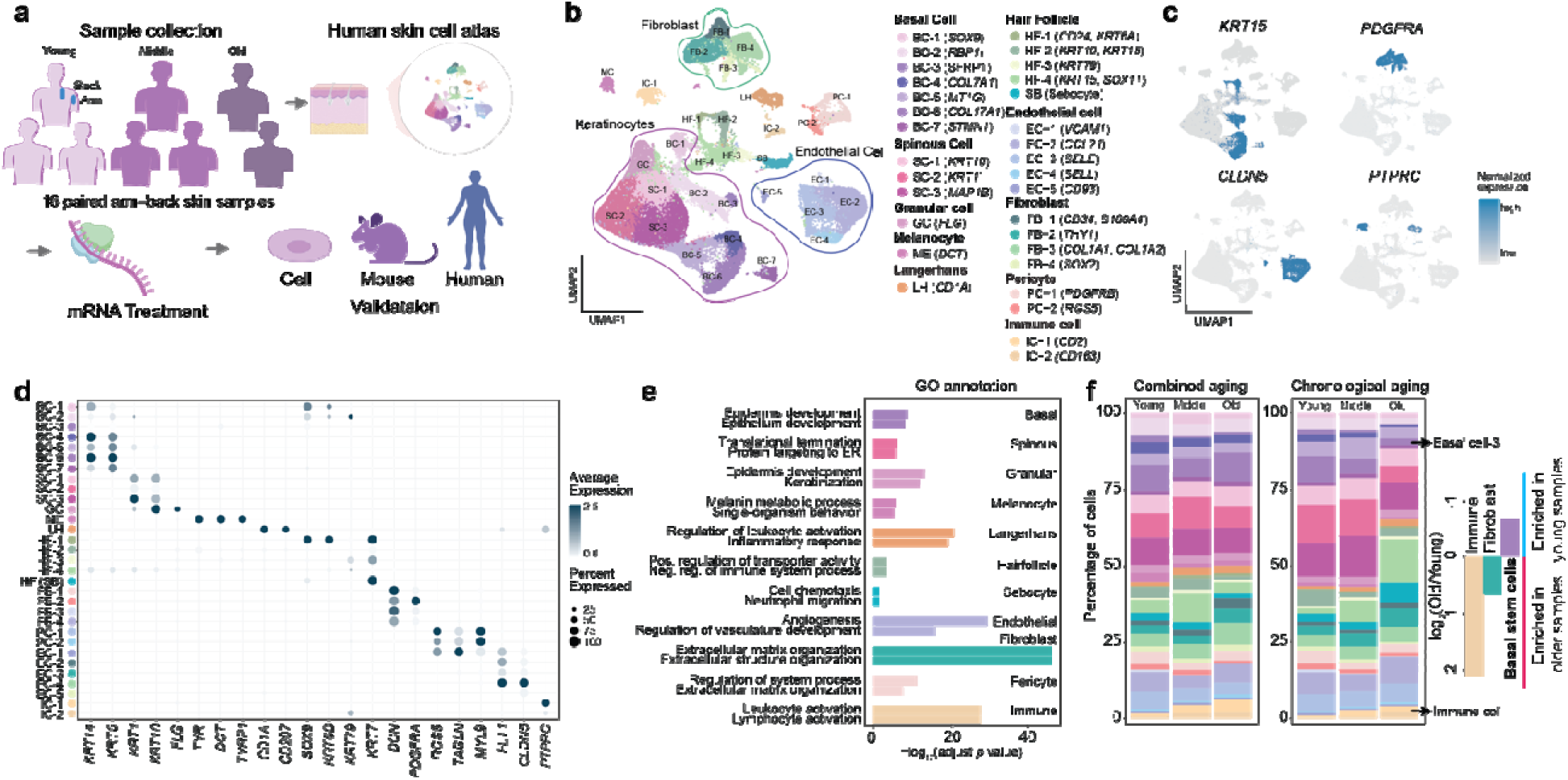
Single-cell characterization of human skin across aging. (a) Overview of the Study. Human skin samples from individuals of varying ages and locations were collected for analysis. Single-cell RNA sequencing (scRNA-seq) was performed to construct a comprehensive cellular atlas. Potential therapeutic targets were subsequently identified from this atlas. The identified targets were validated using small interfering RNA (siRNA) techniques. Following validation, a messenger RNA (mRNA) treatment was developed based on these findings. (b) Uniform manifold approximation and projection (UMAP) plot showing the cell types of human skin. Color denotes the cell type derived from sc-RNA. BC: basal cell; SC: Spinous cell; GC, granular cell; ME: melanocyte; HF, hair follicle; EC, endothelial cell; FB, fibroblast; PC, pericyte; IC, immune cell. (c) UMAP plots show the representative genes of each cell type in human skin. The color key from gray to blue indicates low to high gene expression levels. KRT15, basal stem cells; PDGFRA, fibroblasts; CLDN5, endothelial cell; PTPRC, immune cells. (d) Dot plot showing the expression of representative genes for each cell type. (e) Enriched GO terms for each cell type in combined aging and chronological aging. (f) Bar charts illustrate the proportions of different cell states in human skin across young, middle, and old age groups.

The samples were separated into epidermis and dermis before dissociation to encompass a wide range of cell types, and the cells were characterized using the 10x Genomics 3’ single-cell RNA-sequencing platform. We excluded cells with transcriptomes corresponding to fewer than 200 genes, more than 6000 genes, over 10% mitochondrial gene expression, and doublets, resulting in a total of 109,086 high-quality skin cells. To mitigate batch biases, we integrated data using Seurat ^13,14^ and observed uniform sample distribution in the low-dimensional space (**Extended Data Fig. 1a**).

Subsequently, we identified 31 cell subtypes and 11 cell types through marker-based gene expression on graph-based Leiden clustering. These included all major expected skin cell types: basal cells (BC), spinous cells (SC), granular cells (GC), melanocytes (ME), Langerhans cells (LH), hair follicle cells (HF), endothelial cells (EC), fibroblasts (FB), pericytes (PC), immune cells (IC), and sebocytes (SB). Keratinocytes comprise BC, SC, and GC, with basal stem cells residing in the BC. (**Fig. 1b-d**, and **Supplementary Table 2**). To ensure our analysis was not biased by the stress response associated with tissue dissociation, we removed the known stress response genes, and our 31 cell states remained discernible, suggesting a robust cell characterization ^15^ (**Extended Data Fig. 1b**). Analysis of the top 30 marker genes for each cell type revealed unique transcriptomic features and enriched pathways relevant to their distinct functions (**Fig. 1e, Extended Data Fig. 1c**). For example, basal cell and fibroblasts were enriched for Gene Ontology (GO) terms including epidermal/epithelial development and ECM organization, respectively (**Fig. 1e**). Additionally, genes associated with aging and skin disorders (e.g., squamous and basal cell carcinoma) were mainly identified in melanocytes ^16^. This collective single-cell dataset of human skin provides a foundational resource for studying the processes of skin aging. We also integrated our dataset with all publicly available human skin scRNA-seq datasets (14 datasets in total) and visualized by UMAP (**Extended Data Fig. 1d, Supplementary Table 3**). The integrated public data, comprising 95 donors and 273,178 cells, enables direct cross-study and improved robustness of downstream analyses. These datasets span diverse biological contexts, including healthy skin, wound healing, inflammation, and aging, providing a comprehensive reference framework for robust cross-study comparison. Cell type annotations derived from the public reference datasets showed highly concordant and overlapping clustering with the independently assigned cell types from our dataset, confirming the accuracy and robustness of our cell type annotation.

To investigate the molecular mechanisms underlying commonalities and differences between combined aging (chronological and photoaging) and predominantly chronological aging, we initially characterized skin cell composition changes in both types of aging (**Fig. 1f, Extended Data Fig. 1c**). Although both chronological and combined aging exhibit cellular changes, the magnitude and trends of these changes differ between the two aging types. For example, a change shared between both types of aging is the increase of immune cells (IC-1 and IC-2, yellow). This is consistent with the previous observation that aged skin showed chronic, low-grade skin inflammation ^17^.

In particular, the increase in immune cell proportion was more pronounced when comparing the combined aging groups (middle-aged vs. young, and old vs. middle-aged), indicating that photoaging amplifies skin inflammation (**Fig. 1f**). We also investigated the expression of key menopause-associated genes across age groups in our single-cell dataset, from collected female samples. NRIP1 and KLF9, two regulators of the estrogen signaling and hormone metabolism pathway, were consistently detected and increased with age (**Extended Data Fig. 1e**). This increasing trend is consistent with a gradual decline in estrogen signaling and compensatory activation of downstream transcriptional regulators during skin aging. Furthermore, we observed a reduction in the populations of basal stem cells BC-2, BC-3, BC-4, and BC-6 (characterized by the expression of the KRT15 marker) ^18^. Given that basal stem cells play critical roles in cell adhesion, extracellular matrix organization, cell proliferation, and cellular development and differentiation (**Fig. 1c, Extended Data Fig. 1f**), the depletion of these cells in aged skin suggests a diminished capacity for cell renewal during the aging process ^18^. By conducting pathway analysis on the basal stem cells, we found that combined aging was enriched for pathways involved in cellular migration and skin development; in contrast, chronological aging was enriched for pathways involving morphogenesis and cellular regulation (**Extended Data Fig. 1g, h**). Together, these findings suggest that important genes in identified pathways are involved in the two aging processes and highlight the necessity for targeted therapeutic strategies that address the specific genetic and cellular alterations associated with both types of aging.

### Gene expression programs underlying photoaging and combined aging

To investigate the genetic basis of combined aging and chronological aging, we calculated the differentially expressed genes identified from transcriptomic analysis (DEGs, *t*-test, Bonferroni adjusted *p*-value) for each aging type (**Fig. 2a, 2b, Method**). Most cell types (7 out of 9) exhibited a greater number of DEGs in combined aging compared to chronological aging. This observation aligns with the expectation that combined aging encompasses both photoaging and chronological aging. Notably, in HF and SB, DEGs are mainly overlaps between combined aging and photoaging, suggesting a similarity in the aging processes of these cell types, indicating that these genes are activated by both chronological aging and photoaging.

**Fig. 2.**
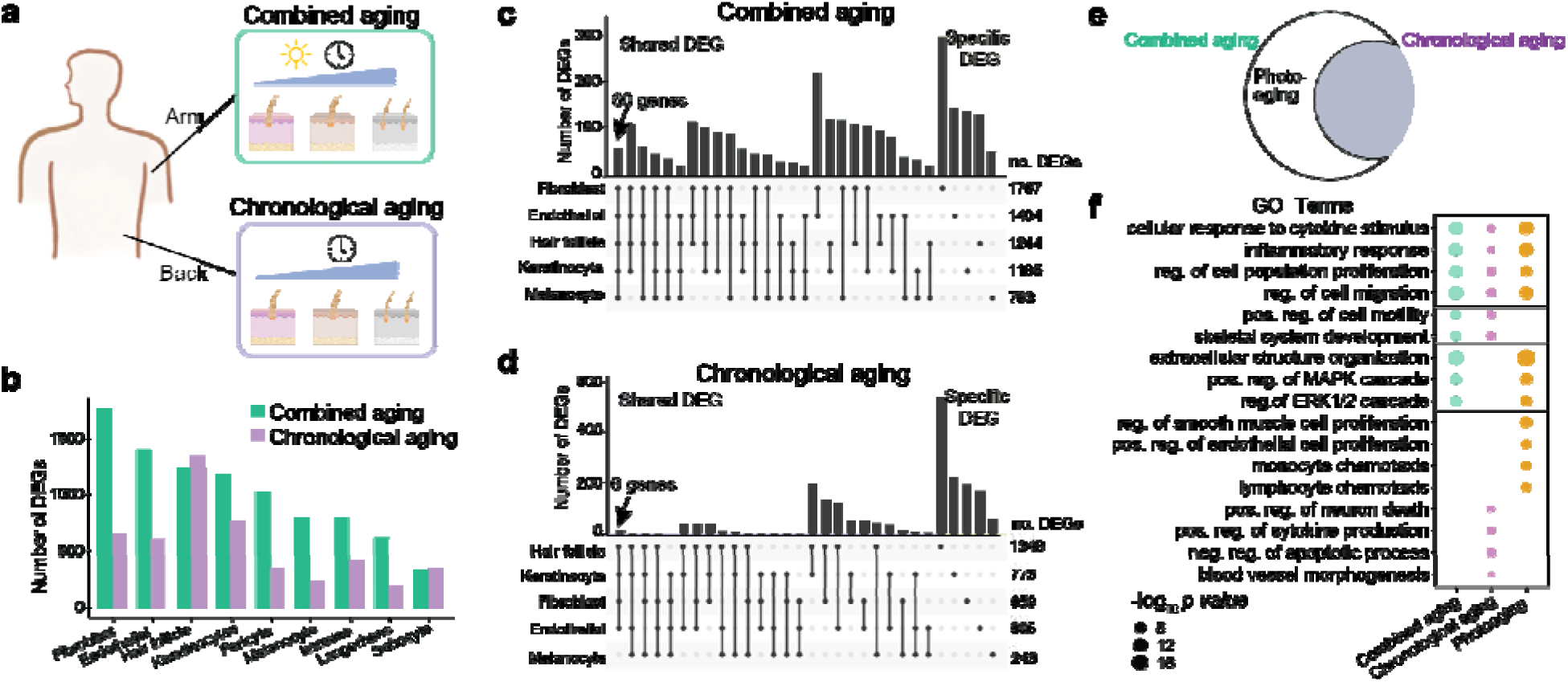
Cellular changes associated with combined aging, chronological aging, and photoaging. (a) Illustration of combined aging (photoaging and chronological aging) and chronological aging. Combined aging is assessed using arm tissues from young, middle-aged, and old individuals, while chronological aging is assessed using back tissues from young, middle-aged, and old individuals. (b) Bar plot depicting the number of differentially expressed genes (DEGs) in various cell types during combined aging and chronological aging. (c) Number of cell type-shared DEGs and cell-specific DEGs in different cell types during combined aging. (d) Number of cell type-shared DEGs and cell-specific DEGs in different cell types during chronological aging. (e) Venn diagram illustrating DEGs across combined aging, chronological aging, and photoaging, with circle size proportional to the number of genes. (f) Top enriched Gene Ontology (GO) Biological Process terms from enrichment analysis of genes selected by combined aging and chronological aging.

Given the differences in DEG numbers, we examined the overlap between these genes. Specifically, we analyzed cell-type-specific DEGs and shared DEGs across five predominant cell types: FB, EC, HF, KC, and ME, which are highly abundant and well-established in skin structure and function. Interestingly, we observed a significant number of cell-type-shared DEGs in combined aging, whereas only a few cell-type-shared DEGs were present in chronological aging. However, the number of cell-type-specific DEGs was similar for both combined aging and chronological aging (**Fig. 2c, d**). This suggests that the differences between combined and chronological aging, specifically photoaging, are primarily attributable to the cell-type-shared genes. We reason that photoaging induces multiple responses in the majority of cell types, as evidenced by the overlapping DEGs across various cell types.

Furthermore, we carefully investigated the shared DEGs across 5 major cell types in combined aging and chronological aging (**Extended Data Fig. 2a, b**). The DEGs exhibited differences in combined aging compared to chronological aging. Specifically, in combined aging, *PRMT9* expression was decreased in endothelial, fibroblast, and melanocyte populations (**Extended Data Fig. 2a**). This aligns with previous studies that indicate a loss of *PRMT9* mRNA expression is associated with cellular senescence and aging in transgenic mice ^19^. Intriguingly, *MEF2A* expression displayed consistent U-shaped trends with aging across all cell types, which may be related to *MEF2A*’s role in cellular proliferative capacity ^20^. *MEF2A* expression increases during growth periods, followed by a gradual reduction during cellular senescence and aging. In contrast, in chronological aging, these same genes demonstrated trends different from combined aging in gene expression over time (**Extended Data Fig. 2b**). These findings underscore the complexity of gene expression dynamics during aging and set the stage for further exploration of their mechanistic underpinnings.

To further elucidate the function of cell-type shared DEGs, we utilized pathway enrichment analysis (**Fig. 2e)**. Specifically, the cell-type shared genes involved in photoaging excluded genes associated with chronological aging from the pool of genes identified in combined aging. Notably, the pathways linked to combined aging closely resemble those of chronological aging but differ significantly from those associated with photoaging. For instance, photoaging is linked more strongly to cellular dysfunction, such as monocyte chemotaxis, while combined aging shows a greater association with skeletal system development and extracellular structure organization (**Fig. 2f**). Subsequently, we examined the enrichment of DEGs related to aging, DNA repair, hallmark EMT, and NFKB (inflammatory) pathways in combined aging relative to chronological aging across various cell types (**Fig 2f, Extended Data Fig. 2c∼f**). Importantly, all cell states demonstrated enrichment of DNA repair-related gene signatures in combined aging (**Fig. 2f, Fig. Extended Data Fig. 2d**), particularly within the middle-aged donor, underscoring the contribution of photoaging to DNA damage ^16^.

Additionally, we evaluated the relative transcriptomic changes of each cellular state across age groups using aggregate measures of DEG fold change (Methods). The “aggregated score” for each cell state was calculated by summing the log2 fold changes of all DEGs within each cell state. Our analysis revealed that the cell types exhibiting the most substantial changes were basal cells (**Extended Data Fig. 2g**). Basal cells are tightly correlated with skin development **(Extended Data Fig. 1e)** and vary in function between chronological and combined aging **(Extended Data Fig. 1f),** indicating their critical role in maintaining skin homeostasis and contributing to the overall process of skin aging.

### Human epidermal keratinocytes act as central regulators of skin aging

Using the analysis of alterations in cell proportions and examination of DEGs, basal keratinocyte cells appear to exhibit a significant association in the aging process. Here, we investigated the heterogeneous characteristics of human epidermal keratinocytes (**Fig. 3a**), with a focus on the basal stem cells. Our dataset provided a comprehensive categorization of the keratinocytes used in this study, which unveiled discrete subpopulations within the stratified epidermis. These subpopulations encompassed seven epidermal basal cells (BC1-7), three spinous subpopulations (SC1–3), and a terminally differentiated cell population (GC) (**Fig. 3b**). To validate our label for basal epidermal subpopulations, we mapped them to a reference dataset on epidermis^21^. Cross-dataset alignment showed highly consistent clustering of basal, spinous, granular, and melanocyte populations across datasets (**Extended Data Fig. 3a**), confirming conserved transcriptional programs and supporting the robustness of our annotation. In particular, our Basal cell-4 and Basal cell-6 clusters mapped precisely to the basal stem cells (BAS-III populations)^21^, confirming the accuracy and robustness of our basal cell clustering. Three SC subtypes demonstrate elevated expression levels of genes associated with barrier function and cell-cell adhesion, including *KLK7* and *DSC1*. Among the seven subgroups of basal cells, our study successfully captured substantial basal stem cells—BC2, BC3, BC4, and BC6, confirmed through the marker gene *KRT15* (**Fig. 1c and Fig. 3d**). In basal stem cells, stemness marker *COL17A1* and inflammatory response genes (*S100A9* and *S100A8*) were highly expressed. In addition, *KRT5*, *POSTN* (Periostin), and *KRT14*, which are related to wound-healing processes, were also enriched in basal stem cells. These basal stem cells offer a promising avenue for investigating skin rejuvenation via stem cell differentiation.

**Fig. 3.**
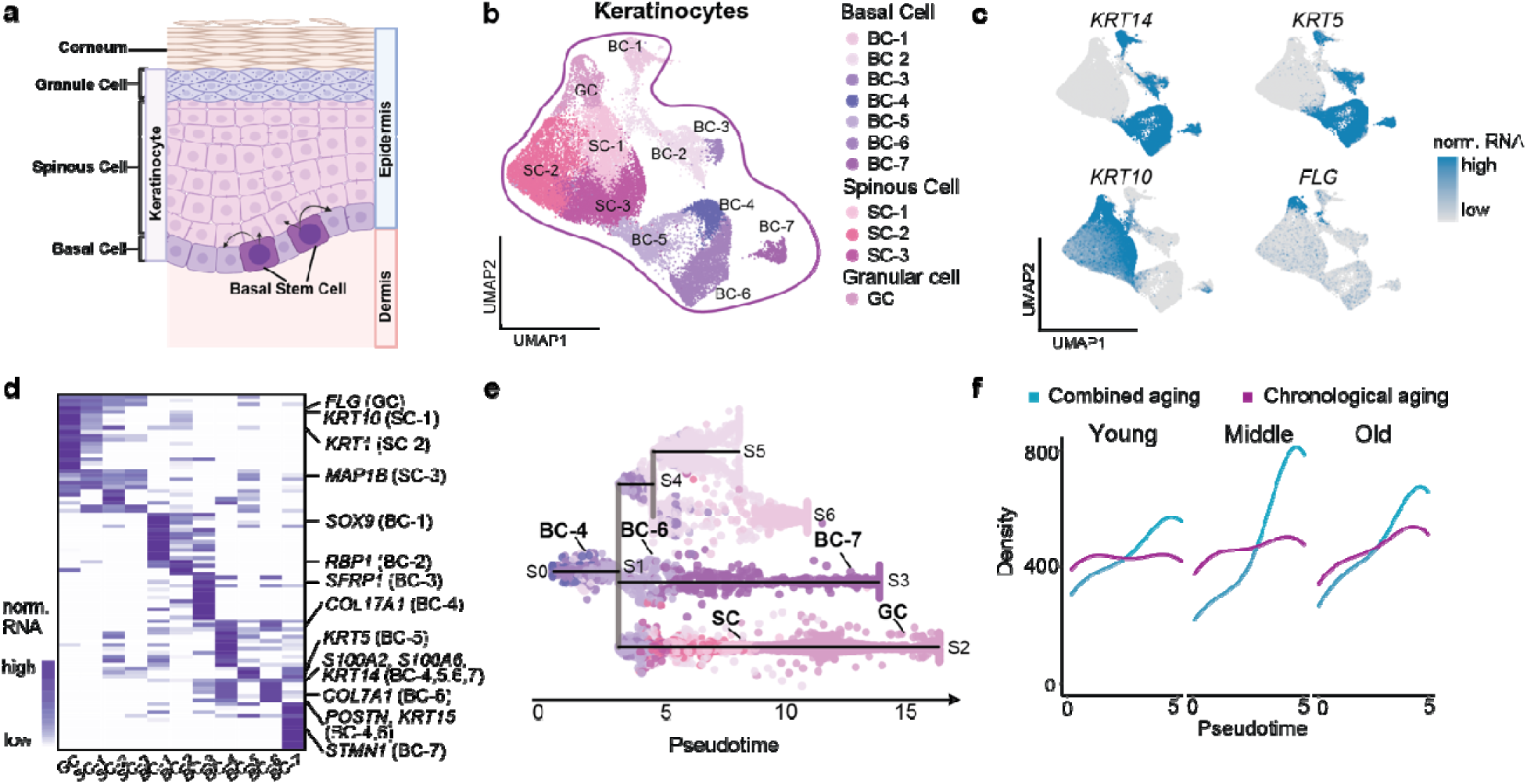
Single-cell transcriptomic classification and differentiation of human epidermal keratinocytes. (a) Schematic representation of human epidermal architecture. Illustration of the epidermal layers, including cornified, granular, spinous, and basal cells. Basal stem cells at the dermal-epidermal junction generate keratinocytes that differentiate upward toward the skin surface, with the dermis providing underlying structural support. (b) UMAP visualizations reveal the categorization of human keratinocytes into 11 subtypes. Basal cells (BC), spinous cells (SC), and granule cells (GC) are indicated based on the expression matrix and their specific markers. Each cell type is distinguished by a unique color. (c) Expression profiles of specified marker genes are depicted across different subtypes. Gradations from gray to blue denote varying levels of gene expression, whereas deeper blue signifies higher expression. Granular cells (GC) are characterized by FLG expression, spinous cells (SC) by KRT10, and basal cell subtype 2 (BC2) by KRT5 and KRT14. (d) Heatmap presenting the scaled expression levels of genes prominently expressed exclusively within all BC, SC, and GC subtypes. The color spectrum, progressing from white to blue, signifies low to elevated gene expression levels. (e) Visualization of the lineage development of keratinocytes along differentiation trajectories. (f) Pseudo-time distribution of cells within lineage S0-S1-S3 for both young and old samples, separated by combined aging and chronological aging scenarios.

Subsequently, pseudotime analysis was performed to elucidate the response of keratinocyte subtypes during skin development (**Fig. 3e**). We identified four distinct cell lineages within epidermal keratinocytes. One lineage, consistent with the epidermal development, progressed from node S0 to S1 to S2, a differentiation trajectory from BC to SC to GC. The remaining three lineages illustrate differentiation within basal cells, specifically from quiescent basal cells to proliferating basal cells. The S0-S1-S3 lineage, which encompasses the differentiation of basal stem cells (BC4 and BC6) to BC7 cells, exhibited significant differences between combined aging and chronological aging (**Fig. 3f**). This lineage demonstrated a gradual deceleration of cell development with age, indicative of reduced basal stem cell renewal capacity during middle and old age. Moreover, middle-aged samples in the combined aging group showed a more pronounced decrement in basal stem cell renewal, highlighting the significant impact of photodamage on basal stem cells during middle age (**Fig. 3f**). Additionally, we explored the S0-S1-S3 trajectory concerning genes involved in the GenAge, the benchmark database of genes related to aging (**Extended Data Fig. 3b**) ^22^. Together, our analysis revealed a consistent pattern wherein these genes measured exhibited decreased expression associated with basal stem cell renewal in aged samples, highlighting their role in regulating basal cell functions across aging contexts.

We also investigated the subtypes of HF, EC, and FB (**Extended Data Fig. 3d-g**), which revealed that gene expression patterns changed the aging process. Consistent with previous studies identifying transcriptionally distinct fibroblast populations in human skin ^23,24^, our single-cell data identified four major fibroblast populations with distinct transcriptional and functional profiles (**Extended Data Fig. 3f**). Papillary fibroblasts (APCDD1⁺, WIF1⁺, COL23A1⁺) situated near the epidermal-dermal junction and characterized by Wnt-pathway regulation and basement-membrane organization; Reticular fibroblasts (DPP4⁺, SFRP2⁺, CTHRC1⁺) enriched in extracellular matrix synthesis and remodeling; Inflammatory fibroblasts (CXCL12⁺, C3⁺, HLA-DRA⁺, ISG15⁺) with cytokine, complement, and antigen-presentation signatures; and myofibroblast (ACTA2⁺, TAGLN⁺, MYL9⁺) involved in contractility, collagen cross-linking, and tissue remodeling. These refined subtypes delineate the structural, inflammatory, and reparative axes of dermal fibroblast heterogeneity in human skin. Analysis of fibroblast subtypes revealed a selective decline in papillary fibroblast numbers and signature genes with age, accompanied by a relative enrichment of reticular and inflammatory fibroblast states, consistent with the compartmental remodeling of the dermis during photoaging^24^ (**Extended Data Fig. 3g**). However, these changes were not as dramatic as the changes observed in basal stem cells, indicating that alterations in these genes may contribute to the reduced renewal capacity observed in aging and that basal stem cells may be an effective target for therapeutic inventions.

### ATF3 mRNA treatment development targeting keratinocytes for skin aging reversal

To identify regulators of skin aging in keratinocytes, particularly basal stem cells, we carefully examined DEGs in keratinocytes (**Fig. 4a**). First, genes were categorized into distinct modules based on their expression patterns across different ages in both combined and chronological aging (**Methods**). We identified two gene modules among eight modules: one characterized by up-regulated genes and the other by down-regulated genes during aging (**Fig. 4b**). Through mapping these two gene modules onto the skin atlas described in **Fig. 1**, we observed the cell type specified in each module. The up-regulated gene module was specific to granular cells (keratinocytes in the stratum granulosum layer), aligning with the role of granular cells in epidermal turnover and the increased propensity for keratosis with age. Our results align with the roles of granular cells in epidermal turnover and the increased propensity for keratinization with age ^25,26^ (**Fig. 4c**). The down-regulated gene module is associated with basal stem cells, suggesting that this module is implicated in stem cell depletion (**Fig. 4c**). To further elucidate the genes in the two modules, we conducted a gene-gene interaction network analysis (**Fig. 4c**) ^27^. In the up-regulated gene network, key genes include *JUP*, *PERP*, *DSC2*, and *DSC3*. These genes are associated with skin integrity and are crucial for cell-cell adhesion. Dysfunction in these genes has been shown to lead to skin fragility ^28^. In the down-regulated gene network, the key genes are *TXNRD1*, *KLF6*, and *ATF3*. TXNRD1 is vital for selenium metabolism and UV protection ^29^. KLF6 is related to cell proliferation and has been reported as a skin-aging gene ^30^. Notably, ATF3, a transcription factor involved in skin development and stress response, was the hub gene of the downregulated gene module, indicating that ATF3 is a critical age-related transcription factor in skin aging (**Fig. 4c, Extended Data Fig. 4a**). The consistent age-associated decline of ATF3 expression in keratinocytes was further confirmed by analysis of the integrated human skin scRNA-seq atlas (**Extended Data Fig. 4b, Supplementary Table 3**). Linear regression analysis showed a clear negative correlation between ATF3 transcript levels and donor age, indicating progressive downregulation of ATF3 during aging in keratinocytes. This trend was reproducible across diverse datasets, sequencing platforms, and anatomical sites, confirming the robustness of the observation. We further performed an in-depth analysis of the dataset spanning fetal, neonatal, and adult human skin^31^ and confirmed a distinct age-dependent decline in ATF3 expression – high levels during fetal stages, a sharp reduction in neonatal skin, and persistently low expression throughout adulthood (**Extended Data Fig. 4c**). These results identify ATF3 as a conserved, age-associated transcriptional marker in human keratinocytes, suggesting a potential role in maintaining epidermal homeostasis and cellular stress responses during aging. To confirm the spatial distribution of ATF3 RNA expression, we examined publicly available spatial transcriptomic datasets across anatomical sites in healthy human skin. We compared the spatial distribution of basal keratinocyte populations (annotated in the original study) to spatial ATF3 mRNA expression patterns (**Extended Data Fig. 4d**) ^32,33^. ATF3 signals were enriched within the regenerative basal niches, consistent with its role in promoting cellular rejuvenation and tissue repair during skin aging. Thus, we hypothesized that ATF3 is a candidate gene for therapeutic targeting in skin aging.

**Fig 4.**
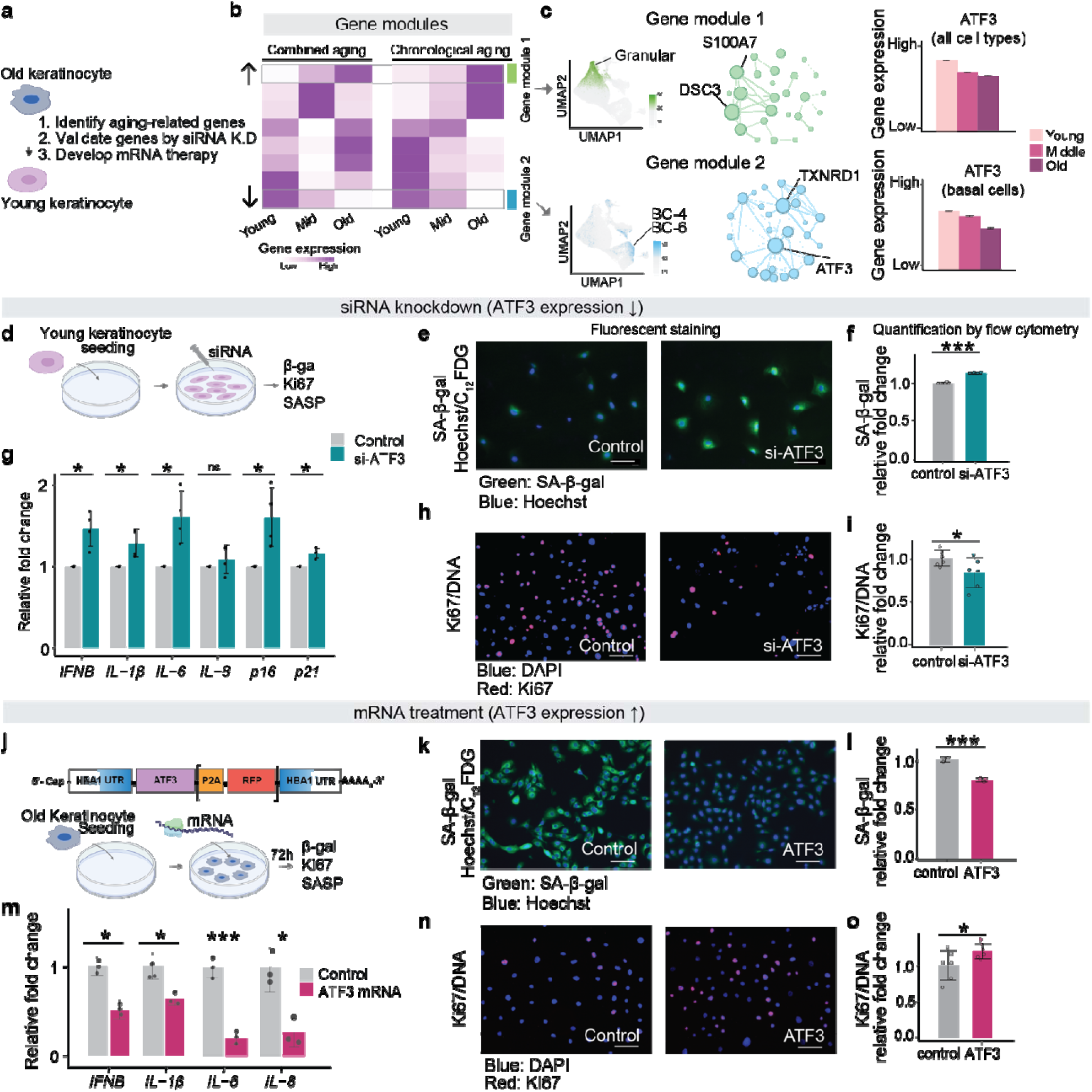
The development of mRNA treatment for human skin rejuvenation. (a) Overview summarizing the development of mRNA therapy for skin rejuvenation. (b) Identification of two distinct “gene modules” emerging during the aging process based on single-cell gene expression clustering: one characterized by upregulated genes and the other by downregulated genes. Genes within these modules are spatially mapped in the skin atlas and integrated into a gene-gene network. (c) Visualization of gene expression patterns within the identified modules using Uniform Manifold Approximation and Projection (UMAP). The gene-gene interaction networks display connectivity, with node size reflecting the number of connections. Bar plots depict expression changes of hub genes across aging stages. (d) Workflow illustrating siRNA-mediated knockdown of ATF3. (e, f) C₁₂-FDG SA-β-gal fluorescence and flow-cytometric quantification in human epidermal keratinocytes after siRNA-mediated ATF3 knockdown. Scale bar, 125 µm. Data are presented as mean ± SEM. n = 3 per group. **p < 0.01. (g) Quantitative PCR (qPCR) analysis of Senescence-associated Secretory Phenotype (SASP) markers in human epidermal keratinocytes upon siRNA-mediated ATF3 knockdown. Data are presented as mean ± SEM. (h, i) Immunofluorescence staining and flow-cytometric quantification of Ki67 in human epidermal keratinocytes following siRNA-mediated ATF3 knockdown. Scale bar, 125 µm. Data are presented as mean ± SEM. n = 6 per group. *p < 0.05. (j) Workflow depicting mRNA treatment by overexpressing the ATF3 gene, designed with in-house untranslated regions. (k, l) C₁₂-FDG SA-β-gal fluorescence and flow-cytometric quantification of human epidermal keratinocytes following mRNA treatment of ATF3. Scale bar, 125 µm. Data are presented as mean ± SEM. n = 3 per group. ***p < 0.001.(m). qPCR analysis of SASP markers in human epidermal keratinocytes upon mRNA treatment of ATF3. Data are presented as mean ± SEM. (n, o) Immunofluorescence staining and flow-cytometric quantification of Ki67 in human epidermal keratinocytes following mRNA treatment of ATF3. Scale bar, 125 µm. Data are presented as mean ± SEM. n = 6 per group. *p < 0.05.

Next, we conducted a siRNA knockdown study in primary human epidermal keratinocytes to validate the role of ATF3 in epidermal aging (**Fig. 4d, Extended Data Fig. 4f**). ATF3 knockdown increased SA-β-gal activity by approximately 20–30% (p < 0.0001) compared with the control group (**Fig. 4e-f**), indicating enhanced senescence. Consistently, ATF3 knockdown reduced the proportion of Ki67-positive proliferating cells by 25% (Ki67/DNA ratio decreased from 1.0 to 0.75) (**Fig. 4h-i**). The knockdown consistently increased the expression of senescence-associated markers by approximately 1.3- to 1.8-fold, including senescence-associated secretory phenotype (SASP) cytokines (IFNB, IL-1β, IL-6, IL-8) and senescence-associated cell-cycle arrest markers (p16 and p21) (**Fig. 4g**). The coordinated up-regulation across multiple senescence-associated genes indicates a biologically meaningful activation of the senescence-associated inflammatory program, rather than random transcriptional fluctuation. Together, these quantitative results and single-cell transcriptomic data support that ATF3 coordinates senescence suppression, epidermal renewal, and collagen production, establishing its role as a central regulatory role as a key age-related gene.

Because we discovered ATF3 expression declines with human skin aging, we leveraged precise and targeted mRNA-based delivery to restore ATF3 expression. We developed a novel mRNA-based therapeutic approach for the potential reversal of human skin aging through upregulation of ATF3 expression using an *in vitro* mRNA delivery method (**Fig. 4j**). The mRNA constructs were designed to include the endogenous 5 and 3 untranslated regions (UTRs) of the HBA1 gene (human alpha-globin-1) to ensure efficient translation and stability (**Supplementary Table 4)**. Following the 5 UTR, a custom human Kozak sequence was inserted, followed by the sequential integration of the ATF3 sequence, a human P2A sequence, a red fluorescent protein (RFP) sequence, an HBA1 3 UTR, and an approximately 107-nucleotide poly-A tail (**Fig. 4j**). These mRNAs were transfected into keratinocytes to validate treatment efficacy. We assessed parameters such as the SASP, cell proliferation, and cellular senescence. After 96 hours of treatment in keratinocytes, a significant reduction in the SASP was observed. Specifically, IL6 levels decreased by 10% to 29% (*t*-test, *p* < 0.0001), and IL8 levels decreased by 12% to 34% (*t*-test, *p* < 0.01). Notably, there was a marked increase in cell proliferation, ranging from 20% to 25% (*t*-test, *p* < 0.01), and a significant decrease in cellular senescence, with reductions between 22% and 25% (*t*-test, *p* < 0.0001). These results indicate the potential efficacy of the ATF3 mRNA in modulating key cellular processes associated with skin aging and senescence, highlighting their therapeutic promise for mRNA-mediated anti-aging interventions (**Fig. 4k-o, Extended Data Fig. 4g-j**).

### ATF3 mRNA treatment restores fibroblast function via keratinocyte-fibroblast communication

Given that the fibroblast is another principal cell type associated with skin aging, we subsequently investigated the effects of delivered ATF3 mRNA in fibroblasts. To determine whether ATF3 mRNA directly reduces senescence in dermal fibroblasts, we transfected primary human fibroblasts with ATF3 mRNA and assessed SA-β-gal activity measured SA-β-gal activity as a senescence marker. The change was not statistically significant. This result suggests that ATF3’s rejuvenating effects are more prominent in keratinocytes, and its impact on fibroblast senescence may occur indirectly through keratinocyte-derived signals.

Consequently, we investigated the intercellular connections between keratinocytes and fibroblasts, motivated by several previous studies^34,35^. By analyzing the cell-cell communication, we identified 62 significant ligand-receptor interaction pairs between keratinocytes and fibroblasts (**Supplementary Table 5, Fig. 5a**). This strong communication was observed in both combined and chronological aging (**Extended Data Fig. 5a, 5b**). Remarkably, the most prominent interactions of ligand-receptor pairs are COL17A1/COL7A1 with the α1β1-complex, α11β1-complex, and α2β1-complex (**Fig. 5b, Extended Data Fig. 5c**). These integrins are key mediators of collagen synthesis and extracellular matrix organization. Consistent with prior studies, we observed a progressive decline in COL17A1 expression with aging (**Fig. 5c, Extended Data Fig. 5d**), a phenomenon previously linked to impaired collagen regulation during skin aging ^36^. Furthermore, ATF3-mediated regulation of the secreted protein CXCL2 serves a pivotal role in mediating the intricate communication between keratinocytes and fibroblasts. ATF3-mediated CXCL2 signaling orchestrates a complex signaling cascade that regulates various aspects of fibroblast behavior, including proliferation and extracellular matrix remodeling (**Fig. 5d)**^37^. These findings provide evidence for intercellular communication mediated by signals, including ATF3, originating from keratinocytes, which influence collagen production in fibroblasts.

**Fig. 5.**
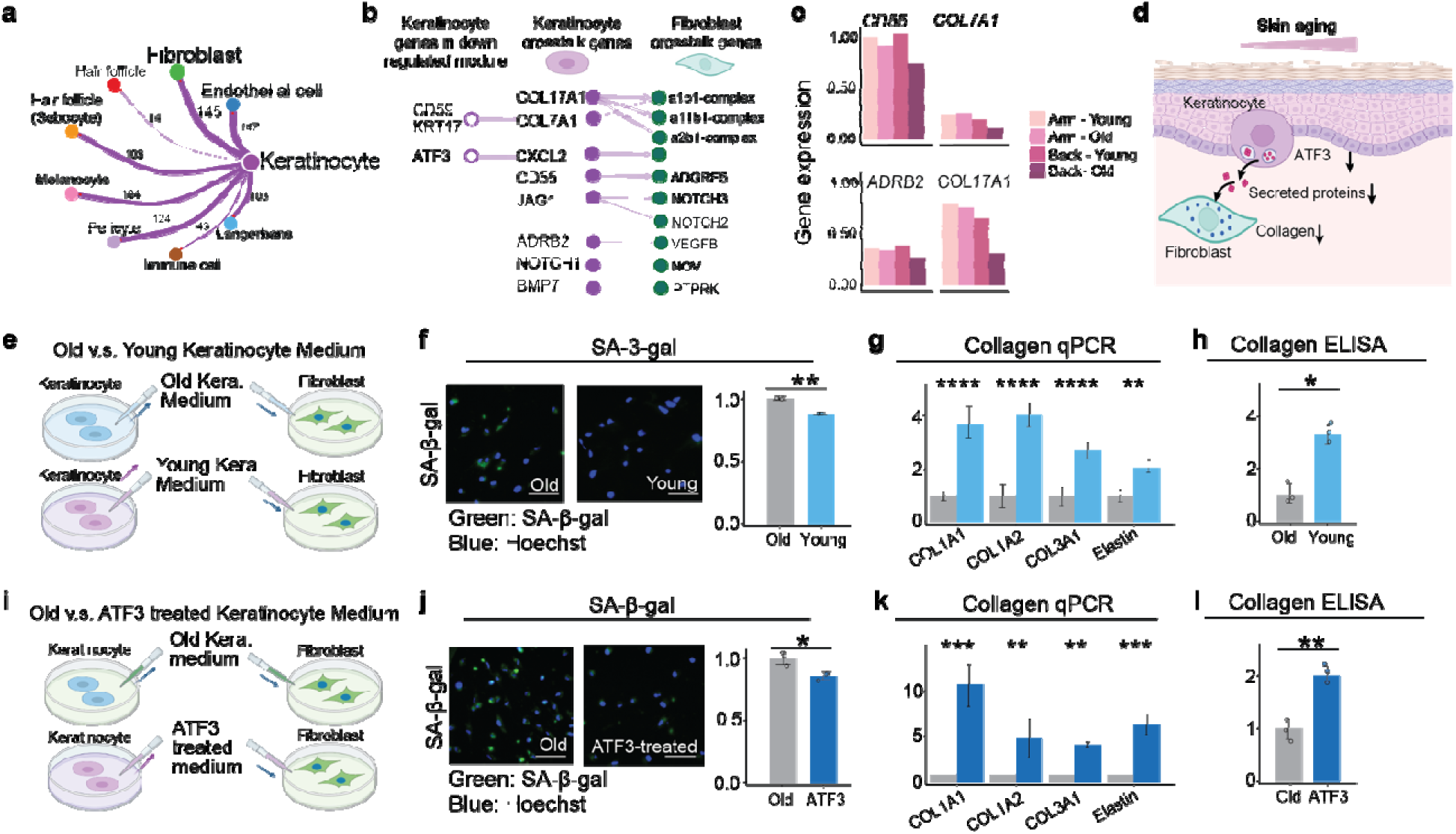
ATF3 mRNA treatment rejuvenates fibroblasts by fibroblast-keratinocyte communication. (a) Network representation illustrating the communication between different cell types, with edge width proportional to the number of identified ligand-receptor pairs. Numbers on edges indicate significant ligand-receptor interactions between cell populations. (b) Significant ligand-receptor pairs contributing to signaling from keratinocytes to fibroblasts were assessed using one-sided permutation tests (*p*-values). (c) Expression profiles of secreted proteins from keratinocytes involved in signaling to fibroblasts. (d) Mechanistic insights into skin aging mediated by secreted proteins from keratinocytes to fibroblasts. (e, f, g, h) Experimental setup involving culture of fibroblasts with conditioned medium from young and old keratinocytes. Evaluation includes Ki67 staining, senescence staining, qPCR of collagen expression, and ELISA of collagen in fibroblasts. Scale bar, 125 µm. Data are presented as mean ± SEM. n = 3 per group. \**p* < 0.05, \*\**p* < 0.01, ****p < 0.0001. (i, j, k, l) Experimental setup involving culture of fibroblasts with conditioned medium from control and ATF3 treated keratinocytes. Evaluation includes Ki67 staining, senescence staining, qPCR of collagen, and ELISA of collagen in fibroblasts. Scale bar, 125 µm. Data are presented as mean ± SEM. n = 3 per group. \**p* < 0.05, \*\**p* < 0.01, ***p < 0.001.

To further characterize these effects, we performed experiments to validate the keratinocyte-mediated communication that influenced fibroblast aging (**Fig. 5e-h, Extended Data Fig. 5e**). Keratinocyte conditioned medium (KCM) collected from old and young primary human basal keratinocytes (largely basal stem cells, **Methods**)^38^ was applied to cultured primary fibroblasts. Both basal keratinocytes and fibroblasts used in our study were isolated from healthy donor skin samples. Treatment with young KCM led to a reduction of approximately 20% fibroblast senescence, as determined by SA-β-gal staining. This treatment also markedly increased the expression of collagen genes in fibroblasts, including COL1A1, COL1A2, and COL3A1, by 2- to 4-fold, and enhanced collagen protein production by approximately 3.5-fold (**Fig. 5d**). These findings demonstrate that young keratinocytes have the capability to promote a youthful state in fibroblasts by enhancing collagen production, thus restoring a regenerative extracellular matrix environment. Furthermore, the keratinocyte-fibroblast communication experiments were independently replicated across samples from six donors, demonstrating the robustness of our findings.

Building upon the findings, we examined whether ATF3 mRNA-treated aged basal keratinocytes could recapitulate the rejuvenating effects of young basal keratinocytes on fibroblasts (**Fig. 5i, Extended Data Fig. 5e**). Encouragingly, KCM transfected with ATF3 mRNA via MessengerMAX significantly reduced fibroblast senescence (**Fig. 5j**). Notably, ATF3 treatment induced a substantial upregulation of collagen and elastin gene expression, resulting in over a 10-fold increase in COL1A1 expression, a 5-fold increase in COL1A2, a 4-fold increase in COL3A1, and a 6-fold increase in elastin expression in fibroblasts (**Fig. 5k∼l**). This aging-associated gene, ATF3, regulates keratinocyte functional integrity, thereby modulating fibroblast senescence and collagen synthesis. These findings support the therapeutic promise of delivering ATF3 mRNA for skin rejuvenation by augmenting collagen synthesis in fibroblasts and mitigating senescence in both fibroblasts and keratinocytes (**Fig. 5a**).

### ATF3 mRNA treatment rejuvenates aging skin in *ex vivo* human skin and *in vivo* mouse models

To evaluate the rejuvenation potential of ATF3 mRNA in aged human skin, we employed an *ex vivo* human skin microneedle delivery model (**Fig. 6a**). Human skin explants provide a highly physiologically relevant system that preserves native tissue architecture, including epidermal and dermal compartments, resident cell populations, and extracellular matrix (ECM) organization, and have been widely applied to assess skin barrier repair and epidermal differentiation ^39^. We obtained full-thickness skin samples from a 62-year-old female donor and topically treated samples with mCherry mRNA (negative control), platelet-rich plasma (PRP, positive control), or ATF3 mRNA, followed by a 96-hour incubation. PRP was included as a positive control given its validated regenerative effects in human skin^40^. For efficient transdermal delivery, mRNA was encapsulated in lipid-based nanoparticles (LNPs) and administered using a microneedle array patch (MAP) ^41,42^, enabling effective skin penetration and localized RNA release into epidermal and dermal cells. Histological and immunofluorescent analyses were then performed to evaluate epidermal renewal (KRT15 and p63) and dermal ECM remodeling (total collagen, elastin, and COL3A1). Immunofluorescence staining confirmed efficient ATF3 mRNA delivery and its functional activity, as evidenced by increased ATF3 protein levels, along with increased basal layer thickness and reduced IFNB protein levels (**Fig. 6b**). The observed reduction in IFNB protein levels further supports ATF3-mediated suppression of SASP at the protein level.

**Fig. 6.**
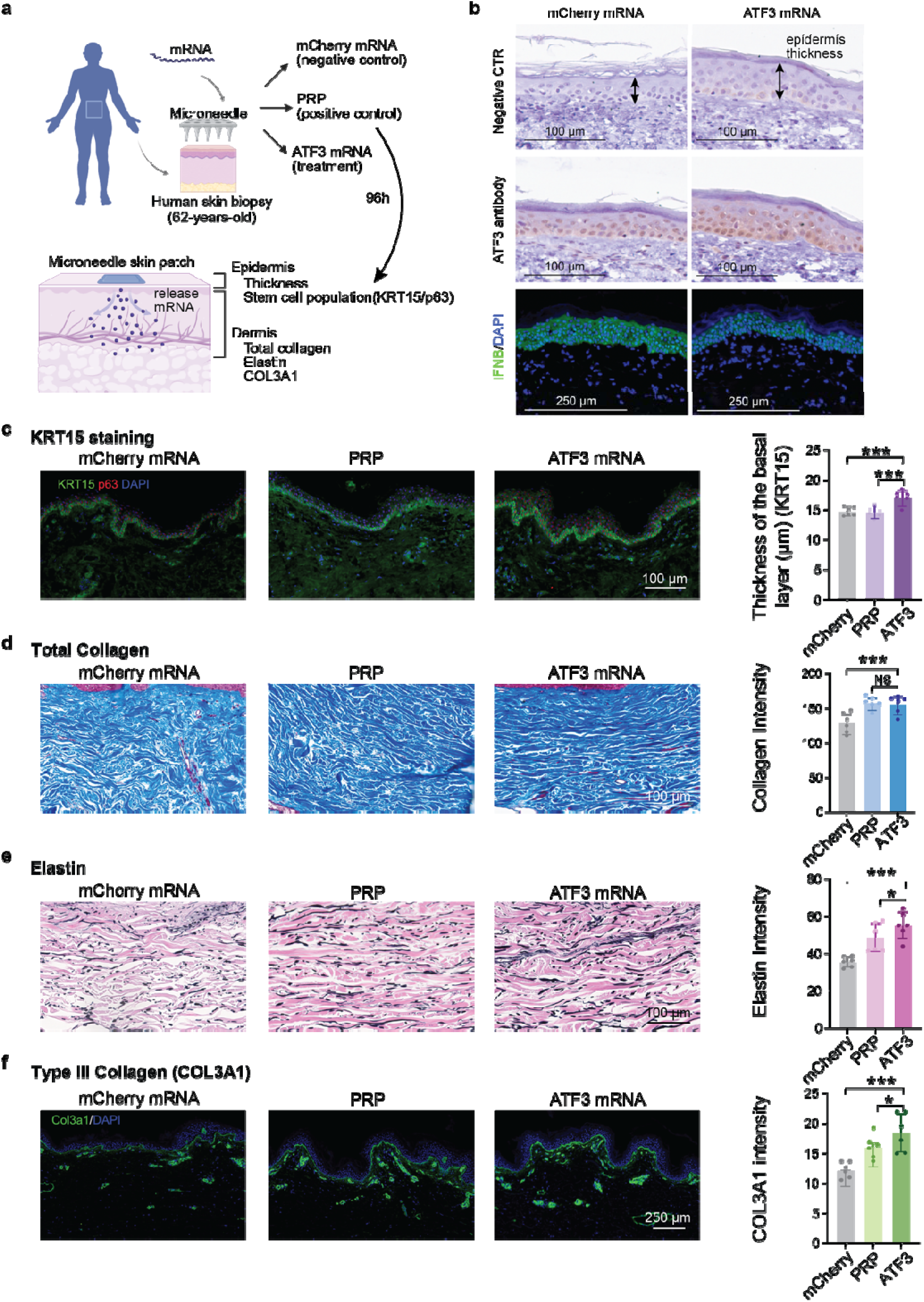
ATF3 mRNA rejuvenates aged human skin *ex vivo* by promoting epidermal renewal and extracellular matrix restoration. (a) Schematic overview of the experimental design showing microneedle-assisted delivery of lipid nanoparticle–encapsulated ATF3 mRNA or control RFP mRNA into 62-year-old human skin punches cultured *ex vivo* for 96 h. Platelet-rich plasma (PRP) was used as a positive control. (b) ATF3 mRNA treatment increases epidermal thickness, reduces IFNB protein expression in human skin explants. Representative H&E, ATF3 immunostaining, and IFNB immunostaining images comparing mCherry control and ATF3 mRNA treatment. (c) Representative histological and immunofluorescent images of epidermal thickness and basal stem cell markers (KRT15 and p63) following treatment with RFP mRNA, PRP, or ATF3 mRNA. Scale bars, 100 µm. (d, e, f) Representative images of dermal extracellular matrix components, including total collagen (Masson’s trichrome), elastin, and type III collagen (COL3A1), across treatment groups. Quantification of basal layer thickness (KRT15+ region), total collagen, elastin, and COL3A1 fluorescence intensity (mean ± s.e.m.). Statistical significance was determined using one-way ANOVA with Tukey’s post hoc test. ***P < 0.001; **P < 0.01; *P < 0.05; NS, not significant.

Remarkably, ATF3 mRNA induced robust rejuvenation effects in aged human skin across all measured parameters. Importantly, ATF3 mRNA treatment markedly increased KRT15 expression, a well-established marker of basal stem cells associated with epidermal stem cell maintenance and regenerative capacity, in the basal layer relative to both mCherry mRNA and PRP groups (**Fig. 6c**). Quantification showed an approximately 1.8-fold increase in basal layer thickness (p < 0.001), indicative of enhanced keratinocyte renewal and activation of the epidermal stem-cell pool. Co-staining with p63, a key transcription factor and marker of epidermal progenitor and stem cells required for basal cell identity and self-renewal, further confirmed the expansion of the progenitor cell compartment within the basal layer.

In the dermis, total collagen deposition markedly increased after ATF3 mRNA delivery, whereas PRP elicited only a modest effect (**Fig. 6d**). Quantitative analysis confirmed a significant increase in collagen intensity in the ATF3 mRNA-treated group (p < 0.001). Similarly, elastin staining revealed a substantial enhancement in elastin fiber density compared with PRP control (p < 0.05) and mCherry control (p < 0.001) (**Fig. 6e**), suggesting that ATF3 mRNA reactivates dermal matrix synthesis and elasticity programs that are typically suppressed with age^43^. Furthermore, immunofluorescence for COL3A1, a key marker of youthful ECM^44^, showed markedly elevated expression in ATF3 mRNA-treated samples compared with both RFP and mCherry groups (p < 0.001) (**Fig. 6f**). COL3A1 is a hallmark of youthful and regenerative dermal extracellular matrix, and its age-associated decline contributes to ECM stiffening and impaired skin regeneration. The selective upregulation of COL3A1 indicates that ATF3 promotes ECM remodeling toward a more regenerative and youthful composition. The *ex vivo* human skin study was conducted three times using skin samples from donors aged 58, 61, and 62 years, yielding consistent results (**Extended Data Fig. 6**).

We next evaluated the rejuvenation potential of ATF3 mRNA *in vivo* using a murine wound-healing model. Because aged skin exhibits delayed repair compared with young skin^45^, we investigated whether ATF3 mRNA could accelerate wound closure and attenuate scar formation. Circular full-thickness excisional wounds were created on the dorsal skin of each mouse, followed by subdermal injection of lipid nanoparticle-encapsulated ATF3 mRNA adjacent to the wound margins^46^. Mice were randomly assigned to either the ATF3 mRNA-treated or control mRNA-treated group (encoding red fluorescent protein, mCherry). ATF3 mRNA treatment reduced scar formation compared with control treatment, indicating enhanced regenerative repair *in vivo* (**Extended Data Fig. 6**).

Together, these findings demonstrate that ATF3 mRNA delivery rejuvenates aged human skin, enhancing epidermal stem-cell renewal and dermal matrix reconstruction beyond the regenerative effects achieved by conventional PRP treatment. These results position ATF3 as a central regulator of skin regeneration and a promising therapeutic candidate for reversing age-associated decline in skin regenerative capacity.

## DISCUSSION

Aging is a dynamic biological reality and a complex phenomenon ^47^. Like other organs, human skin experiences aging in two forms, chronological and environmental, particularly from photoaging. In this study, we uncovered the complex mechanisms behind both types of skin aging and developed an mRNA treatment for skin rejuvenation. Central to our research was a pioneering single-cell atlas that enabled us to map the genes responsible for specific cellular processes involved in skin aging and the identification of the ATF3 gene as a crucial player in the skin aging process. Previous studies have linked ATF3 to carcinoma, epidermal differentiation, and cellular stress responses ^48–50^. In contrast, our data identify a distinct role for ATF3 as a rejuvenation and regeneration-associated factor in aging human skin, acting primarily within basal stem cells. This distinction likely reflects differences in experimental scope and analytical resolution. By profiling a large expanse of human skin at single-cell resolution, our study captures a broad spectrum of cell types, with extensive coverage of keratinocyte subtypes, enabling refined analysis of ATF3 activities. Our finding reveals an unrecognized function of ATF3 in maintaining stem cell competence and tissue repair capacity during aging, representing a conceptual advance in understanding its role in skin biology and the molecular regulation of aging.

We have pioneered the creation of a comprehensive human skin aging atlas of photoaging and chronological aging. This resource provides the aging field with an unprecedented view of the varying cellular states across the lifespan under both UV-exposed and non-exposed conditions. Our study contributes to the understanding of how chronological aging and photoaging may involve distinct pathways. This aligns with other studies that link aging with inflammatory responses and cellular dysfunction ^51^. Of note, our findings reveal convergent pathways and genes that are influenced by both chronological and photoaging, enabling the development of more effective anti-aging strategies. Previous studies on aging have typically concentrated either on photoaging, examining genes impacted by ultraviolet radiation, or on chronological aging, resulting in an incomplete understanding of the aging process. Our research offers a unique framework for investigating both internal and external factors contributing to aging, providing significant insights for skin aging and broader aging research.

By leveraging the understanding of both photoaging and chronological aging, we revealed the pivotal role of basal stem cells in skin aging. Previously, studies have focused on fibroblasts as the primary mediators of age-related changes ^24,52^, but our findings demonstrate that basal stem cells undergo far more dramatic transformations than fibroblasts. These results significantly shift the current understanding of cellular contributors to skin aging and suggest that basal stem cells are key targets for therapeutic interventions.

Furthermore, our study elucidates a significant mechanism of intercellular communication between keratinocytes (particularly the basal stem cell subset) and fibroblasts. ATF3 mRNA treatment reduced the gene expression of senescence markers and robustly enhanced the collagen and elastin production in fibroblasts. This highlighted the pivotal role of fibroblast-keratinocyte communication in driving functional rejuvenation of the dermal niche, emphasizing the complexity of skin aging. It also indicates that key regulatory genes in one cell type, such as ATF3, influence both epidermal and dermal layers. Understanding intercellular communication broadens the potential for developing therapeutic interventions for aging in various organs, as well as for addressing other skin disorders, including inflammatory conditions and alopecia.

Finally, our research highlights the promise of a first-of-its-kind mRNA therapy for skin rejuvenation. mRNA technology has been successfully employed in vaccines, demonstrating its safety and efficacy in human populations ^53,54^. This modality is cost-effective and presents a promising alternative for anti-aging therapies and other medical applications such as wound healing. The use of mRNA is particularly significant due to its ability to induce precise and targeted gene expression, which can directly address molecular mechanisms. Furthermore, mRNA-based approaches do not result in permanent genomic alterations, in contrast to genome-editing technologies. Instead, this treatment is controlled easily with dosing regimens, and the delivered mRNA is temporarily expressed in treated cells. mRNA therapies also offer rapid development timelines, allowing for swift adaptation in response to emerging skin-related conditions. Our study demonstrates that ATF3 mRNA delivery produces robust rejuvenation in aged human skin by simultaneously restoring epidermal stem-cell function and reactivating dermal extracellular-matrix remodeling. The improved performance of ATF3 mRNA compared with platelet-rich plasma illustrates the therapeutic relevance of selectively targeting stem-cell regulatory pathways rather than relying on broad, nonspecific regenerative interventions. Furthermore, the reduced scarring observed *in vivo* supports ATF3 mRNA as a promising regenerative therapy with potential clinical applicability for reversing age-related skin decline and enhancing tissue repair.

Given the growing interest in regenerative medicine, the potential of mRNA to enhance skin rejuvenation and improve dermal integrity represents a promising and feasible treatment approach. Our study of skin rejuvenation and innovative technology could revolutionize strategies for combating aging in other organs and other disorders, paving the way for more effective therapies, and ushering in a new era of targeted interventions with far-reaching implications for human health and longevity.

## METHODS

### Ethics statement and tissue acquisition

This study on skin aging involved the participation of individuals who graciously donated samples of their healthy adult skin. Before their involvement, these donors provided written informed consent, ensuring ethical compliance with the No.3 Zhongshan Hospital Research Ethics Committee (REC reference: 08/H0906/95+5). To ensure consistency and relevance, strict criteria were applied for the selection of donors. None of the individuals had diabetes, were obese, or suffered from other skin diseases. Their body mass index (BMI) fell within the range of 23 to 28, establishing a healthy baseline for the research.

The samples collected for analysis were of dimensions 10 mm x 10 mm, and meticulous efforts were made to procure them from identical positions on the arm and back of each donor. This standardized approach allows for accurate comparisons between the three age groups: young individuals (age 23), middle-aged individuals (age 53), and elderly individuals (age 85) (**Supplementary Table 1)**.

### Generation of single-cell suspension and Single-cell RNA-seq

To prepare a single-cell suspension from adult skin, healthy skin samples were first cut into thin strips while immersed in phosphate-buffered saline (PBS). The top 200 μm layer was isolated using a dermatome with a Pilling Wecprep blade and a 0.008-gauge Goulian guard. Grid slits were introduced into the skin sheets for enzymatic access, followed by a 1-hour treatment with 2 U/ml dispase II in RPMI at 37°C. Subsequently, the epidermis was peeled from the dermis, and both fragments were separately digested in a petri dish at 37°C 5% CO_2_ in RF-10 media with 1.6 mg/ml type IV collagenase (Worthington, CLS-4) for 12 hours. RF-10 media consists of Roswell Park Memorial Institute media (RPMI) (Sigma, R0883) supplemented with 10% (v/v) fetal calf serum (FCS, Life Technologies, 10270106), 100U/ml Penicillin (Sigma, P0781), 100 µg/ml Streptomycin (Sigma, P0781) and 1% (v/v) L-Glutamine (Sigma, G7513). The work was conducted within class II biological safety cabinets using autoclave-sterilized equipment. The media was collected using a serological pipette and filtered through a sterile 100 μm cell strainer (BD Falcon, 352360). After washing the petri dish and strainer with RF-10 media to gather any remaining cells, centrifugation was employed to pellet the cells at 500g for 5 minutes. The supernatant was discarded and the pellet was gently resuspended in 1 ml RF-10 media through pipetting up and down. Cell counting was performed using a hemocytometer after staining 10 μl of the sample with an equal amount of 0.4% trypan blue (Sigma, T8154) to identify dead cells.

Single cells were captured using droplet emulsions, and scRNA-seq libraries were generated following the manufacturer’s instructions using the Chromium 10x Single-Cell Instrument (10x Genomics) and 10x Genomics Chromium Single Cell 3’ GEM Library and Gel Bead Kit v2. Each channel was loaded with cells, aiming for 10,000 cells per sample, with cell concentration measured by Moxi GO II (Orflo Technologies). The cDNA was amplified for 12 cycles in the Bio-Rad C1000 Touch Thermal cycler with 96 Deep-Well Reaction Module. Amplified cDNAs and final libraries were assessed using a Fragment Analyzer (AATI) with a High Sensitivity NGS Analysis Kit (Advanced Analytical) to determine the average fragment length of the 10x cDNA libraries. Quantification of the libraries was done via qPCR using the Kapa Library Quantification kit. Subsequently, the libraries were diluted to a final concentration of 2 nM and pooled together for sequencing. All libraries were sequenced using the NovaSeq 6000 Sequencing System (Illumina) with an average coverage of 50,000 raw reads per cell, sequenced in a 28 x 10 x 10 x 90 bp configuration.

### Processing raw data from scRNA-seq

Raw data from single-cell RNA sequencing (scRNA-seq) were processed using the default parameters of the Cell Ranger software suite (10x Genomics). The quality assessment of the sample-specific FASTQ files was performed based on the Cell Ranger counts, which were aligned to the human reference genome (GRCh38) using the STAR aligner to create a gene expression matrix. Each transcript’s expression level was determined by the count of unique molecular identifiers (UMIs) assigned to that transcript. The gene expression matrices were utilized for downstream analyses.

### scRNA-seq data analysis and cell-type identification

Filtered feature barcode matrices from Cell Ranger ARC were analyzed using the Seurat R package (v4.3.0) ^13^. Following the initial Cell Ranger metric assessment, cells with transcriptomes corresponding to fewer than 200 genes or more than 6,000 genes and more than 10% of mitochondrial genes were excluded from downstream analyses. After quality control, a total of 61,106 cells remained and were utilized for subsequent bioinformatic analyses. Sequencing reads for each gene were normalized and scaled using the SCTransform function in Seurat. We performed data integration using Seurat to mitigate potential batch effects across samples and experiments ^13^. Subsequently, we applied dimensionality reduction and clustering techniques to analyze the integrated data. Cell clustering was performed using the ‘FindClusters’ function with a resolution of 0.8, utilizing the first 30 PCs to define cell identities. Dimensionality reduction was performed using the ‘RunUMAP’ function, and the results were visualized with Uniform Manifold Approximation and Projection (UMAP). Marker genes for each cluster were determined using the Wilcoxon rank-sum test with the ‘FindAllMarkers’ function. Only genes with log fold change > 0.25 and adjusted *p* value < 0.05 were considered as marker genes. The identified marker genes for each cluster are presented in **Supplementary Table 2**.

### Identification of aging-associated differentially expressed genes (DEGs)

To identify aging-associated DEGs between the old and young groups (O/Y), middle-aged and young groups (M/Y), and old and middle-aged groups (O/M) for each cell type, we utilized the “FindMarkers” function in Seurat. The log fold change and adjusted p-value for each DEG were calculated using the non-parametric two-sided Wilcoxon rank-sum test. Only DEGs with an absolute average log fold change greater than 0.25 and an adjusted *p* value less than 0.05 were considered significant.

### Gene Ontology (GO) Analysis and KEGG pathway analysis

GO and KEGG pathway analysis of DEGs was performed by EnrichR (version 3.2) ^55^ and visualized with the ggplot2 R package. Representative terms selected from the top 20 ranked GO terms or KEGG pathways (*p* < 0.01) were displayed.

### Pseudotime analysis

Epidermal cell developmental trajectories were computed using the STREAM algorithm [35]. Gene filtering was performed using the filter_genes function with the parameter min_num_cell set to 5. Variable gene selection was then conducted with the select_variable_genes function, using a loess_frac parameter of 0.01. To initialize the tree structure, the parameter n_cluster was set to 10. The elastic principal graph was obtained with the following parameters: epg_alpha set to 0.015, epg_mu set to 0.2, and epg_lambda set to 0.02.

### Gene set scoring

Individual pathways were collected from the GenAge and KEGG database and scored in single cells Seurat AddModuleScore function and visualized in ggpubr R package (version 0.5.0). Individual pathways were tested for significance using a Wilcoxon rank-sum test and Bonferroni corrected through the ‘rstatix’ R package (version 0.7.2).

### Perturbation plot and pathway enrichment analysis

Pathway activity was determined using the ‘clusterProfiler’ R package (version 3.0.4) using the enrichGO function with default parameters. Enrichment plots were generated using the ‘enrichplot’ (version 3.1.8) ^55^. Differentially expressed genes were determined by the Seurat function FindMarkers with default parameters. The overall ‘perturbation score’ for each cell state was calculated by summing all log_2_ fold changes for DEGs in each cell state.

### Integration of public scRNA-seq datasets

Public human skin scRNA-seq data from 14 studies comprising 93 samples and 85 donors were collected and uniformly processed (**Supplementary Table 3**). Cells passing quality-control filters (nCount_RNA > 100, nFeature_RNA > 200, mitochondrial fraction < 10%, ribosomal fraction < 60%, hemoglobin fraction < 5%) were retained. One low-quality sample with fewer than 100 cells post-QC was excluded, yielding 285,887 cells for downstream analysis.

Data integration was performed using Seurat v5. Expression values were normalized and scaled with regression of nCount_RNA and mitochondrial percentage, followed by PCA. To account for protocol heterogeneity across studies, including separate epidermal and dermal sequencing versus whole-skin dissociation. Harmony was applied on a per-donor basis to correct batch effects while preserving biological variation across anatomical sites. Unsupervised clustering was used to define major cell types based on canonical markers. Epithelial, stromal, and immune compartments were subsequently subclustered at resolutions of 0.6, 0.8, and 1.0, respectively.

### Spatial transcriptomics analysis

Spatial transcriptomics data were obtained from a published study profiling healthy human skin across anatomical sites using 10x Genomics Visium^32^. We analyzed four representatives non-UV-exposed body skin samples. Basal keratinocyte annotations provided in the original dataset were used to visualize their spatial distribution alongside ATF3 expression. Cell populations annotated in scRNAseq data were computationally predicted on global ST sections using cell2location^33^. Predicted cell abundances shown by color gradients per spot in tissue architecture (H&E) images. Gene expression values were scaled to the 99th percentile for visualization, with basal keratinocytes shown in blue and ATF3 expression in green, overlaid on corresponding H&E images.

### Gene modules

Gene modules were clustered by performing a hierarchical cluster analysis using a set of dissimilarities for the gene expressions. Initially, each object is assigned to its cluster and then the algorithm proceeds iteratively, at each stage joining the two most similar clusters, continuing until there is just a single cluster. At each stage, distances between clusters are recomputed by the Lance--Williams dissimilarity update formula according to the particular clustering method being used. Gene-gene interactions within a module were identified by mapping the genes to the STRING network ^27^.

### Cell-cell communication analysis

To assess cell-cell communication molecules between different cell types, we used CellPhoneDB software (version 1.1.0) to infer the intercellular communication network from single-cell transcriptome data ^56^. Only receptors and ligands expressed in more than 10% of the cells in the specific cell types were considered in the analysis. First, by randomly permuting the cluster labels of all cells 1,000 times, we determined the mean of the average receptor expression level of a cluster and the average ligand expression level of the interacting cluster. Then, we performed pairwise comparisons between all cell types and obtained a likelihood of p-value to filter the false-positive interaction. Only interactions with *p* < 0.05 were considered to be significant (**Supplementary Table 5**).

### Cell culture

Human primary keratinocytes were isolated from healthy skin samples obtained during surgical procedures, which were generated as part of the surgical procedure. This process inherently selects for basal epidermal stem cells located in the interfollicular epidermis, as only this population retains the capacity for long-term proliferation in vitro. This selection has been well-documented since the foundational work by Rheinwald and Green^57^, who demonstrated that only basal cells can form proliferative colonies under these conditions. Other keratinocyte lineages, having already committed to differentiation, are incapable of sustained proliferation and therefore cannot survive or expand in culture. This is because terminal differentiation in the epidermis is coupled to cell cycle exit, as shown by Zhu, et.al^38^. Hence, in our system, the term “primary human keratinocytes” are largely basal stem cells of the interfollicular epidermis.

Subcutaneous adipose tissue was excised from the human skin specimens, and the tissue pieces were incubated in dispase solution (Stemcell, 07913) at 4°C overnight. Following digestion, the epidermis was separated from the dermis, chopped into small pieces, and incubated in 0.05% Trypsin-EDTA (Thermo Fisher Scientific, 25300054) for 10 minutes. The human primary keratinocytes were collected through a 70 mm cell strainer, seeded in a precoated dish, and cultured in supplemented Keratinocyte SFM (K-SFM, Thermo Fisher Scientific, 17005042), which included 0.2 ng/mL human recombinant EGF, 30 μg/mL BPE, and 1% antibiotic/antimycotic. The cells were cultured at 37°C under an atmosphere of 5% CO₂ in the air. Cells were cultured in K-SFM supplemented with 30ug/ml bovine pituitary extract (BPE) and 0.3ng/ml recombinant epidermal growth factor (rEGF) with 5% CO_2_ at 37°C. Human dermal Fibroblasts were obtained from the Aging Cell Culture Repository (NIA, Coriell Institute for Medical Studies) and were cultured in DMEM medium containing 10% FBS at 5% CO_2_ at 37°C.

### ATF3 staining in human skin tissue

Immunohistochemistry staining was performed according to previously developed protocols ^58^. Paraffin-embedded sections were deparaffinized with three washes of 100% xylene and rehydrated through a series of graded alcohols (100%, 100%, 100%, 95%, and 80%), followed by a brief wash in distilled water. Antigen retrieval was carried out using heat-mediated treatment in sodium citrate buffer (pH 6.0) for 20 minutes. Sections were then treated with 3% H_2_O_2_ to quench endogenous peroxidase activity, followed by a 1-hour incubation in blocking buffer at room temperature. Primary Anti-ATF3 [CL1685] antibody incubation was performed overnight at 4°C. Following PBS washes, HRP-conjugated secondary antibodies were applied, and detection was conducted using DAB. The sections were counterstained with eosin and dehydrated through a series of graded alcohols (80%, 95%, 95%, 95%, 100%, 100%, and 100%) and 100% xylene before cover-slipping with a resinous mounting medium.

### Validation of ATF3 Expression with Published Data

To validate ATF3 expression variation between young and old samples, we manually curated sequencing data from four research papers (PMID: 33238152, 35069694, 34031030, 32327715), selecting samples from healthy individuals. These samples were stratified into young (<30 years) and old (>60 years) groups according to the criteria set in the original publications. The curated data were integrated using Scanpy version 1.9.2 and annotated based on either the original publications or in-house developed models. ATF3 expression levels were compared by averaging the raw counts for keratinocytes in each sample. Statistical analysis was performed using a Student’s t-test.

### siRNA Treatment

Primary human basal stem cell keratinocytes were plated the day before treatment in a 6-well plate to achieve 70-80% confluency at the time of treatment. 150 pmol of Dharmacon ON-TARGETplus siRNA (Non-targeting: D-001810-01-05; ATF3: J-008663-05-0002) was complexed with 7.5 µL of Lipofectamine RNAiMax (Invitrogen, 13778075) according to the manufacturer’s instructions and then added to the cells. The keratinocytes were incubated with the siRNA complexes for six hours, then the medium was changed. Cell samples were collected at 48 hours post-transfection for RT-qPCR to assess target knockdown, and SASP RT-qPCR was performed on samples collected at 72 hours post-transfection.

### Senescence-associated **β**-galactosidase staining

Senescence-associated β-galactosidase (SA-β-gal) staining was performed using the Senescence β-Galactosidase Staining Kit (Cell Signaling Technology, 9860). Briefly, cells were rinsed with PBS and fixed with 1x fixative solution for 15 minutes. Fresh X-gal stock solution was prepared by adding 1 ml of DMSO to 20 mg of X-gal. The β-Galactosidase staining solution was then prepared according to the manufacturer’s instructions. The fixed cells were rinsed with PBS twice, and a total volume of 1 ml of the staining solution was added to each well. The plate was sealed and incubated in a 37°C dry incubator for 18 hours. The staining results were observed using a microscope, and β-galactosidase-positive cells were considered senescent cells, which were counted in 3 randomly chosen fields. The SA-β-gal staining was further quantified by flow cytometry.

C12FDG staining for SA-β-gal was performed by initially incubating cells with 100 nM Bafilomycin A1 (VWR, 102513) for 1 hour. Subsequently, cells were incubated for an additional hour with 100 nM Bafilomycin A1 and 33 nM 5-Dodecanoylaminofluorescein Di-β-D-Galactopyranoside (C12FDG, ThermoFisher, D2893). Then, flow cytometry was conducted using a BD LSR II Flow Cytometer, and data were analyzed using FlowJo software (Version 10.10.0).

### Flow Cytometry

Cells were initially stained with Hoechst 33342 (Dojindo) for 30 minutes to label the nuclei before dissociated into a single-cell suspension. We used the eBioscience™ Intracellular Fixation & Permeabilization Buffer Set (Invitrogen) for fixation and permeabilization. A total of 1 mL of the fixation working solution was added to each sample, which was then incubated for 30 minutes at 4°C. After incubation, the samples were washed twice with 2 mL of 1X Permeabilization Buffer. Post-washing, the cell pellet was resuspended in 100 µL of 1X Permeabilization Buffer and incubated with 1 µg of Alexa Fluor 488 anti-human Ki-67 Antibody (BioLegend, 350532) for at least 30 minutes at room temperature. Following this incubation, the samples were washed twice with 2 mL of 1X Permeabilization Buffer. The final stained cells were resuspended in 150 µL of Cell Staining Buffer (BioLegend) and analyzed using a BD® LSR II Flow Cytometer.

### Ki67 Immunohistochemistry Staining

For Ki67 immunohistochemistry, cells were stained with Hoechst 33342 (Dojindo) for 30 minutes before fixation in a 24-well plate. After three washes with PBS, cells were fixed with 4% paraformaldehyde (PFA) at room temperature for 20 minutes. Following fixation, the supernatant was removed, and the cells were permeabilized with 0.1% Triton-X at room temperature for 20 minutes. The cells were then washed three additional times with PBS. A 25 µL aliquot of a 1:200 dilution of Alexa Fluor® 647 anti-human Ki-67 Antibody (Abcam, ab281928) was added to each well. Cells were stained overnight at 4°C and subsequently inspected using a FLoid™ Cell Imaging Station.

### Immunohistochemistry Staining for the Target Gene

5 µm-thick sections of formalin-fixed paraffin-embedded (FFPE) human skin specimens from young (20-30 years old) and aged (≥ 80 years) patients were deparaffinized and rehydrated using a graded ethanol series. After quenching endogenous peroxidase activity (BLOXALL Blocking Solution, Vector Laboratories), heat-induced epitope retrieval was performed using citric acid-based antigen unmasking solution (Vector Laboratories) at 110°C for 15 minutes in a programmable antigen retrieval chamber (ARC, Biocare Medical). Subsequently, tissue sections were blocked with 2.5% normal goat serum (Vector Laboratories) at room temperature for 20 minutes, followed by incubation at 4°C overnight with mouse anti-ATF3 (1:100, Abcam ab191513) in a humidified chamber. The next day, after washing with phosphate-buffered saline, tissue sections were incubated with ImmPRESS HRP Goat Anti-Mouse IgG Polymer Reagent (Vector Laboratories) for 30 minutes in a humidified chamber according to the manufacturer’s recommendation. Chromogenic detection was performed using ImmPACT DAB Peroxidase (Vector Laboratories) prepared according to the manufacturer’s instructions and applied to the tissue sections for a total of 7 minutes, followed by counterstaining with Hematoxylin QS (Vector Laboratories), dehydration in 100% isopropanol, and coverslipping with toluene-based mounting medium (VectaMount Express, Vector Laboratories). The following semi-quantitative scoring rubric was employed to evaluate staining: 3+: >50% of cells had strong nuclear/nucleolar staining, 2+: <50% of cells had variable nuclear/nucleolar staining, and 1+: <50% of cells had weak primarily nucleolar staining.

### mRNA synthesis

Double-stranded DNA (dsDNA) template vectors of ATF3 (**Supplementary Table 4**) were PCR amplified from the donor plasmid (pUC-GW-Kan, Genewiz) using primers. The resulting dsDNA template vectors were purified using the QIAquick PCR Purification Kit (Qiagen). Gene block inserts (IDT) were cloned into the dsDNA template vector using Gibson Assembly (New England Biolabs). The resulting circular dsDNA plasmid was then transformed into TOP 10 competent cells (Thermo Fisher Scientific). The cells were cultured for 12 hours at 37°C. A single bacterial colony was picked and cultured for 12 hours in LB at 37°C. The next day, the plasmids were extracted and purified using the QIAprep Spin Miniprep kit (Qiagen). The resulting plasmids were digested at 37° C for 30 minutes using BamHI-HF (New England Biolabs) and subsequently purified using the QIAquick PCR Purification Kit. Linearized dsDNAs from subsequent steps were used to synthesize the mRNAs. mRNAs were *in vitro* transcribed for 12 hours at 37°C using a MEGAscript T7 transcription kit (Thermo Fisher Scientific). The transcription mixture contained 1 µg of template dsDNA, 5 mM of ATP nucleotide-triphosphate, 5 mM of CTP nucleotide-triphosphate, 3 mM of N1-methyl pseudouridine-5-triphosphate (TriLink Biotechnologies), 4 mM of CleanCap reagent AG (TriLink Biotechnologies), 0.1 U Inorganic pyrophosphatase (Thermo Fisher Scientific). Transcribed mRNAs were purified using LiCl precipitation following the MEGAscript T7 transcription kit instructions.

### mRNA treatment

Old (Female 62 years old) primary human epidermal keratinocytes were cultured in the K-SFM, supplemented with 30 μg/ml bovine pituitary extract (BPE, Gibco) and 0.3 ng/ml epidermal growth factor (EGF, Gibco) at 37 °C and 5% O_2_ atmosphere, splitting the cells every 2-4 days to maintain monolayer coverage. Two days before transfection, cells were seeded in 6-well plates at a density of 200,000 cells per well.

For transfection, the purified mRNA was prepared at a concentration of 1 µg/µl in UltraPure Distilled Water (Invitrogen). For each treated well, the mRNA was diluted in 125 µl of Opti-MEM Reduced Serum Medium (Thermo Fisher Scientific) at the desired concentration. In a separate tube, MessengerMAX reagent was diluted in 125 µl of Opti-MEM at a ratio of 2 µl of MessengerMAX per 1 µg of mRNA. The diluted mRNA and MessengerMAX reagent were combined, gently mixed, and incubated at room temperature for 10 minutes to allow complex formation.

After the incubation period, 250 µl of mRNA-MessengerMAX complexes were added dropwise to each well of the cell culture, with gentle swirling to ensure uniform distribution. After transfection, the plate was incubated at 37°C in a CO₂ incubator to promote optimal cellular uptake. After 30 minutes, the transfection medium was removed, and cells were washed twice with PBS. Media was replaced with fresh K-SFM, and cells were returned to 37°C incubator.

### Fibroblast-keratinocyte communication

Primary human epidermal keratinocytes from a 27-year-old donor (PromoCell, C-12003) and 56-year donor (Lifeline Cell Technology, C-0025) were cultured in Keratinocyte Growth Medium (Lifeline Cell Technology, LL-0007), which was devoid of Gentamicin and Amphotericin B but supplemented with 100U/ml Penicillin-Streptomycin. At 80% confluence, the cell was washed twice with Hanks′ Balanced Salt solution (Sigma Aldrich, H6648) and cultured in fresh low-glucose Dulbecco’s modified Eagle’s medium (Gibco, 11885-084) for an additional 48 hours. The culture media were then collected and employed as conditioned media for experiments on the same day. A comparative negative control was established by incubating low-glucose Dulbecco’s modified Eagle’s Medium at 37°C for the same duration.

Human dermal fibroblasts were sourced from a 28-year-old female from the Aging Cell Culture Repository (NIA, Coriell Institute for Medical Studies) and were maintained as monolayer cultures in low-glucose Dulbecco’s modified Eagle’s medium (DMEM, Gibco, 11885-084) supplemented with 10% fetal bovine serum (Gibco, A3160501) and 100 U/ml Penicillin-Streptomycin until they reached 75% confluence. The fibroblasts were then seeded at 1×10^5^ cells/ml and incubated overnight to allow cell attachment. Afterward, the cells underwent two washes with PBS and were subjected to a 24-hour serum starvation period in low-glucose DMEM without fetal bovine serum. This was followed by a 24-hour incubation with a mixture consisting of 50% low-glucose DMEM and 50% keratinocyte-conditioned media. The cell count in each well was then normalized with the Cell Count Normalization kit (Dojindo, C544) and tested for SA-β-gal activity.

### RNA-isolation and real-time reverse transcription-quantitative polymerase chain reaction (qPCR)

Total RNA was extracted using the RNeasy Plus Mini Kit (Qiagen, 74104) according to the manufacturer’s protocol. cDNA synthesis was performed using the SuperScript III First-Strand Synthesis System (Invitrogen,18080051), and qPCR was performed in technical triplicate using the KAPA SYBR FAST Universal qPCR kit (KK4602) on a Roche LightCycler 96 instrument. Relative quantitation was determined using the ΔΔCt method with GAPDH as an endogenous normalization control. The primer sequences for qPCR are listed in **Supplementary Table 6**. qPCR assays were performed using a minimum of three independent samples. Group differences were evaluated using an independent samples t-test. Results are reported as mean ± SEM, p-value < 0.05 is considered statistically significant.

### ELISA

To quantify the total amount of Collagen Type I in the culture, both the cells and the supernatant were collected separately post-treatment. The cells were lysed on ice using RIPA lysis buffer for 20 minutes. Collagen Type I concentrations in both the cell lysate and supernatant were quantified using the Human Collagen Type I ELISA Kit (Abcam, ab285250). The assay was performed by following the manufacturer’s protocol to ensure accuracy and consistency in measurements.

### Preparation and characterization of lipid nanoparticles for mRNA delivery

mRNA-loaded lipid nanoparticles (LNPs) were prepared at an N:P molar ratio of 6:1. To generate the organic phase, 4-(dimethylamino)-butanoic acid (D-Lin-MC3-DMA, MedChemExpress), cholesterol, 1,2-dioleoyl-sn-glycero-3-phosphoethanolamine (DOPE, Avanti), and 1,2-dimyristoyl-rac-glycero-3-methoxypolyethylene glycol-2000 (DMG-PEG 2000, Avanti) were mixed in pure ethanol at a predetermined molar ratio. The aqueous phase was generated by diluting mRNA in 20 mM acetate buffer (pH 4). Lipid nanoparticles were synthesized by mixing the aqueous phase and the organic phase at a volumetric ratio of 3:1 by pipetting or pulse vortexing depending on the preparation scale. The lipid nanoparticles were then dialyzed in phosphate buffered saline for at least 2 h using a Pur-A-LyzerTM dialysis kit (Sigma) at 4°C and concentrated using centrifugal ultrafiltration devices (Amicon, Millipore). Encapsulation efficiency was determined using the Quant-iT Ribogreen RNA assay (Invitrogen) following the manufacturer’s protocol. Size, polydispersity index, and surface charge measurements were obtained by dynamic light scattering and zeta potential measurements using a Nanosizer ZS (Malvern Instruments).

### Preparation of microneedle patches for transdermal mRNA delivery

Microneedle patches were fabricated using a centrifugation method to cast polymers into a polydimethylsiloxane mold as previously described ^42^. Each microneedle patch consisted of an 11 by 11 array with each needle having a radius of 150 µm and a height of 600 µm. To generate the hyaluronic acid backbone polymer, the carboxyl groups of sodium hyaluronate (60kDa, LifeCore Biomedical) were activated using N-(3-dimethylaminopropyl)-N’-ethylcarbodiimide (1:4 molar ratio) and N-hydroxysuccinimide (1:2 molar ratio) at room temperature for 30 minutes, before reaction with cystamine dihydrochloride (1:10 molar ratio) at room temperature for 12 hours. The resultant amino-modified hyaluronic acid was dialyzed in water for one week, then freeze-dried and stored at −20°C until use. For the preparation of the microneedle patch, amino-modified hyaluronic acid (10% w/v) was cast into the mold by centrifugation and then freeze-dried. After removal from the lyophilizer, a second layer consisting of an 8-arm PEG-NHS crosslinker (10% w/v, PEGworks) was deposited on the mold, centrifuged, and again freeze-dried. Next, the drug layer containing either concentrated lipid nanoparticles with 8% w/v sucrose (mCherry or ATF3 microneedle groups) or 8% w/v sucrose only (empty microneedles) was added by centrifugation. Finally, the backing layer was added by dropwise addition of 15% w/v poly (D, L-lactide-co-glycolide) (Resomer RG 505, Sigma). Microneedle patches were allowed to dry at 4°C for at least three days prior to being peeled off the molds and stored at 4°C until further use.

### *In vivo* wound-healing experiments

Aged C57B6/J mice (17-18 months old; female) were anesthetized with isoflurane, and the planned injection area on the dorsal skin was shaved and outlined using a sterile marker. Lipid nanoparticle (LNP)-formulated mRNA encoding RFP (control) or ATF3 was delivered via a 50 µL intradermal injection on 4 days before day relative to wounding (Day −4). Mice in the control group received 12 µg RFP mRNA, and mice in the treatment group received 12 µg ATF3.

Following recovery, mice were monitored daily for clinical observations, including local reaction, weight change, fur texture, mobility, and general behavior. Four days after mRNA administration (Day 0), full-thickness excisional wounds were generated by tenting the skin against a hard, sterile surface and applying a 6-mm biopsy punch to two symmetrical dorsal regions per mouse. Wounds were left uncovered and allowed to heal by secondary intention. Mice received 1mg/kg extended-release buprenorphine (Wildlife Pharmaceuticals) for pain management at least 1 hour before surgical intervention.

Digital images of wounds were obtained daily with a calibrated camera. Wound size was quantified by measuring the longest and shortest wound diameters using digital calipers. Body weight and signs of systemic toxicity were evaluated throughout the study period. At Day 15, wound and peri-wound skin tissues were excised using a 12-mm biopsy punch for downstream histological analysis.

### Human skin explants and treatments

Full-thickness human skin explants (12 mm punch biopsies) were placed dermal side down on transwell inserts (Costar) in contact with growth medium (DMEM, Gibco) supplemented with 10% human serum (Gemini Bio Products) and 1% antibiotic/antimycotic. The epidermal surface of each explant was maintained at the air–liquid interface. Before culture, microneedle application was used to deliver either control RFP or ATF3 mRNA directly into the explants. Platelet-rich plasma (PRP) was added to the growth medium. Treated skin explants were incubated for 72 h, harvested, and fixed overnight in 10% neutral-buffered formalin for paraffin embedding or directly embedded in OCT compound (Tissue-Tek), snap-frozen at –80°C.

### Histology and immunofluorescence of human skin explants

Paraffin sections of fixed skin explants were processed for hematoxylin and eosin (H&E) staining and histological analysis. Masson’s trichrome staining was performed using the VitroView Masson’s Trichrome Stain Kit (VitroVivo Biotech, VB-3016), and elastin staining using the VitroView Verhoeff-Van Gieson Elastin Stain Kit (VitroVivo Biotech, VB-3019).

For immunofluorescence in OCT, sections were air-dried for 15 minutes at room temperature and fixed in 4% paraformaldehyde for 15minutes. The cryosections were permeabilized in 0.5% Triton X-100/PBS for 30 minutes. Sections were blocked and incubated overnight at 4 °C with the following primary antibodies: anti-KRT15 (Biolegend, #833904), anti-p63 (Abcam, ab124762), anti-IFNB (Abcam, ab85803), and anti-Col3a1 (GeneTex, GTX27778). Alexa Fluor-conjugated secondary antibodies were applied for 1 h at room temperature. Nuclei were counterstained with Hoechst 33342 (1 µg/ml, Invitrogen).

For immunohistochemistry, deparaffinized sections were subjected to heat-induced antigen retrieval using citrate-based antigen unmasking solution (Sigma). Endogenous peroxidase activity was quenched with 3% H_2_O_2_ for 5 min, followed by blocking in 2.5% normal serum (Vector Laboratories) for 30 min. Sections were incubated overnight at 4 °C with anti-ATF3 (Sigma Aldrich, HPA001562). A biotinylated secondary antibody (Vector Laboratories) was applied for 1 h. Signal was developed using ImmPACT DAB (Vector Laboratories), and nuclei were counterstained with hematoxylin before dehydration and mounting.

Images were acquired using the NanoZoomer s60 digital scanner (Hamamatsu Corp.). Quantification of fluorescence signals was performed using Fiji/ImageJ software.

## Acknowledgments

We thank members of the Church lab for their critical reading of the manuscript and helpful discussions, including Chun-Ting Wu, Katelyn Buehring, Harlan Stevens, and Michael Somekh. We are grateful to Ricardo Rodriguez Vidal in the Zhang lab for supporting the qPCR experiments and Dr. Yi Hu from the Feinberg Lab for support on the cell culture. We thank Dr. Chenli Liu and SIAT for their support, which contributed to the preliminary RNA sequencing efforts that facilitated this study. We thank the BPF Genomics Core Facility at Harvard Medical School for their expertise and instrument availability that supported this work. We gratefully acknowledge Aaron Tam and Amanda Graveline for their contributions to the *in vivo* mouse experiments. This work was funded by the Leo Foundation (LF-OC-20-000420), the grant from the American Academy of Dermatology (AAD), the Wyss Director Fund, and the Wyss Validation Fund.

## Author contributions

L.L. and G.M.C. conceived the idea of the study. L.L. performed computational analyses, wrote the manuscript, worked with all co-authors, and was involved in all experiments and analyses. R.A. contributed to the computational analyses. P.R. and A.L.J. contributed to the collection of public single-cell datasets. X.P., M.Y., and Z.T. contributed to the mRNA treatment section. E.M., P.H., Z.T., Z.Z., J.L., D.D.M., A.A.T., Y.S.Z., A.M.P., X.L., and M.S. performed the knockdown and communication experiments. S.S., F.B., E.D.C., and A.M. contributed to the primary keratinocyte collection, keratinocyte-fibroblast communication experiments, and *ex vivo* human skin study. M.D., J.M.T., and N.A. contributed to the *in vivo* mouse study and microneedle skin patch. Y.Z. and Y.W. contributed to skin sample collection. G.F.M. and S.M.M. performed the imaging experiments. G.F.M., J.S.M., J.M.T., and T.S.K. contributed to the clinical concepts. L.L., S.M., and G.M.C. supervised the work. All authors proofread the manuscript.

---

**Extended Data Fig. 1.**
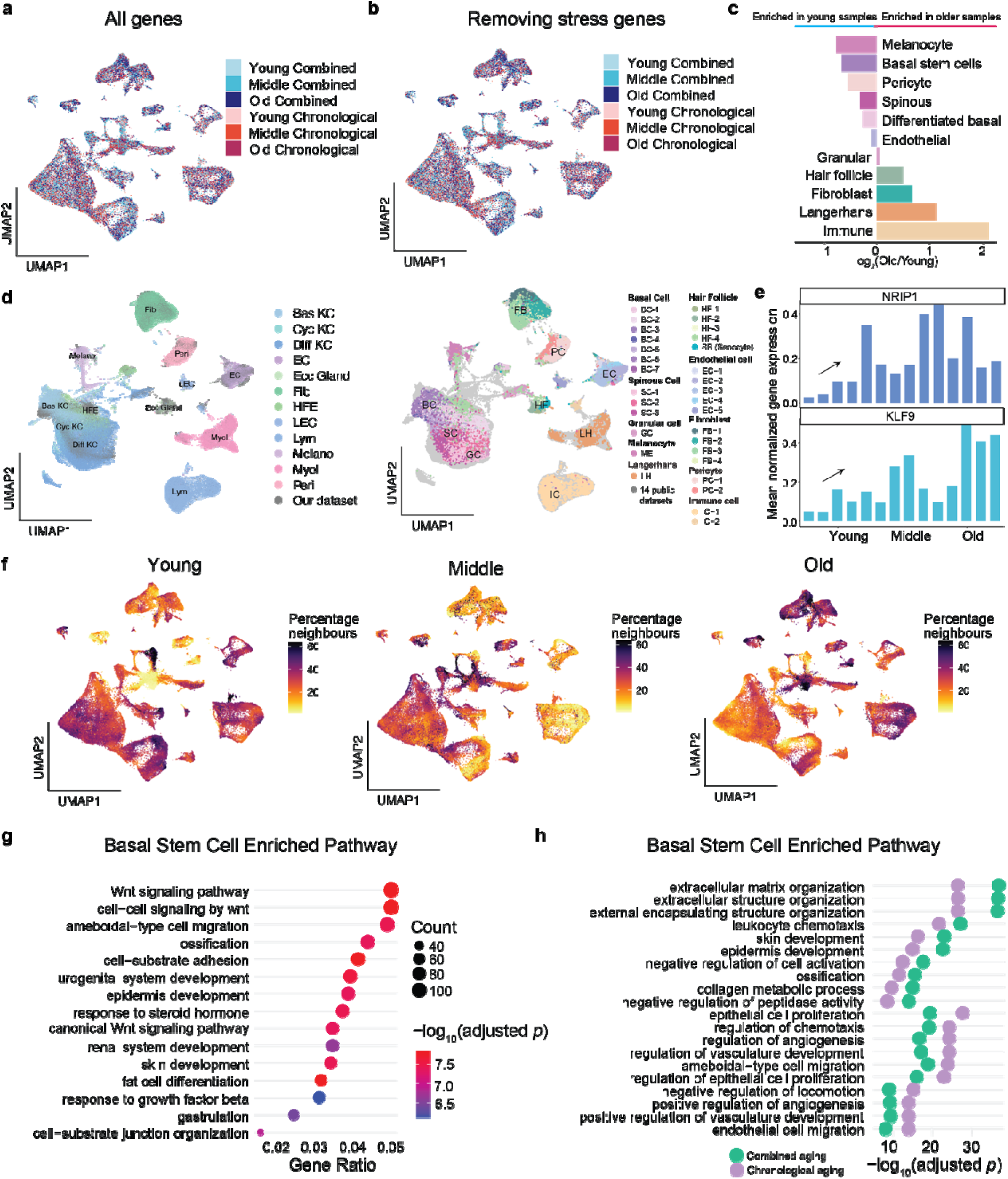
Single-cell alterations associated with skin aging. (a) UMAP plot of all cells of human skin, colored by sample origin. (b) UMAP plot of all cells of human skin by removing the stress genes, colored by sample origin. (c) The cell number variations between young and old samples across all cell types. (d) UMAP of the integrated dataset combining our samples with 14 public scRNA-seq datasets, annotated using reference-derived cell types (left). The same cells colored by our independent annotations (right). (e) Menopause-associated upregulation of NRIP1 and KLF9. Mean normalized expression of NRIP1 and KLF9 across Young, Middle, and Old / Post-menopause skin samples. Both genes show increased expression with age and are highest post-menopause. (f) Expression differences among young, middle-aged, and old samples by measuring the percentage of neighbor cells. (g) Dot plot identifying top pathways upregulated in the basal stem cells. The size of the dots shows the number of genes while the color of the dot indicates adjusted p-value. (h) Pathway enrichment analysis for basal stem cells, with pathways ordered by adjusted p-value and dots colored by aging type. The plot displays the top 10 pathways specific to each aging type.

**Extended Data Fig. 2.**
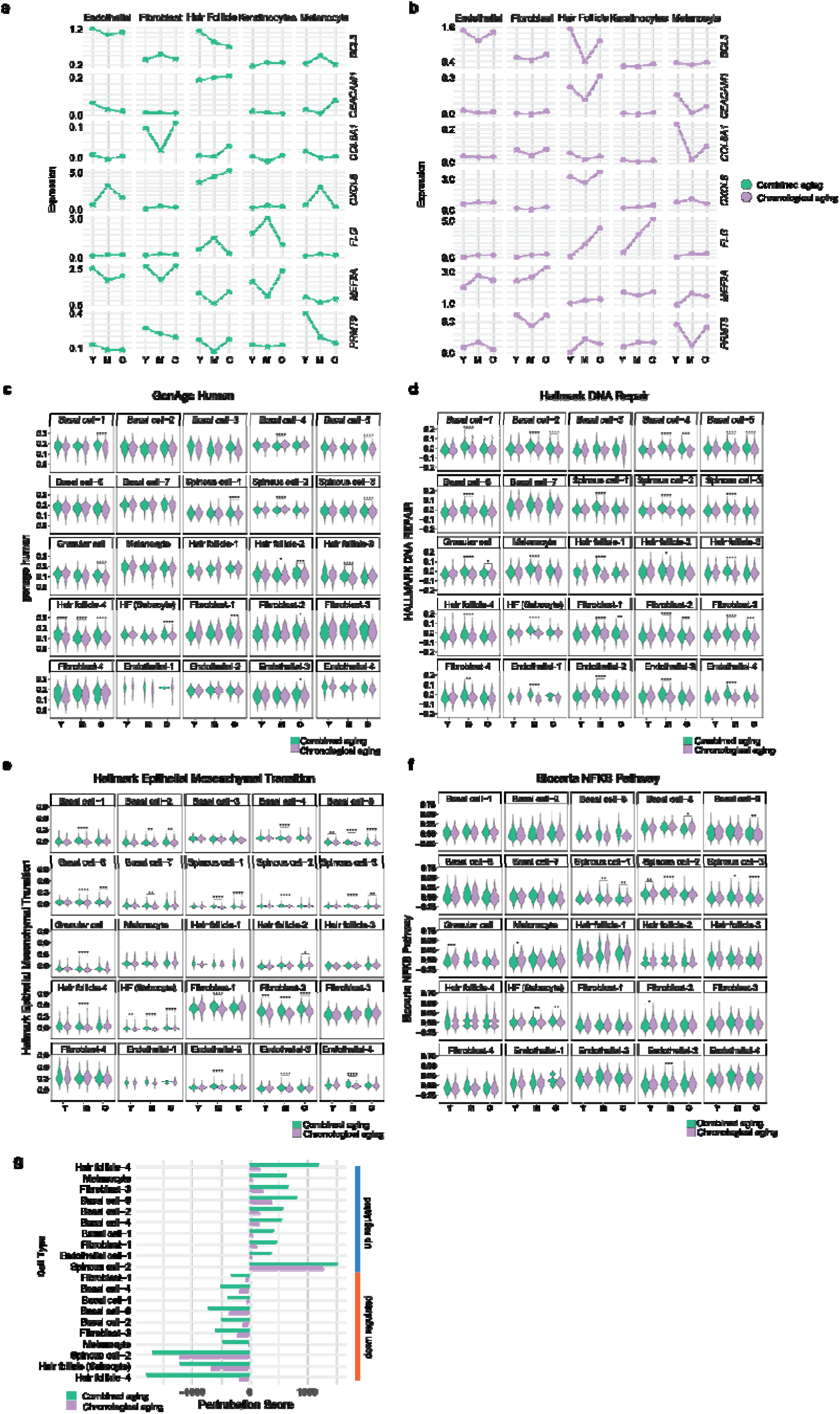
Gene expression dynamics across age during combined and chronological aging. (a) Line plot across ages showing trends in 6 out of 60 cell-type-shared DEGs expression in combined aging. (b) Line plot across ages showing trends in 6 cell-type-shared DEGs expression in chronological aging. (c-f) Line plot of specific DEGs across ages and cell types in combined aging. Gene set scoring for GenAge (d), DNA repair (d), hallmark EMT (e), and NFKB (f) related genes across cell states. Violin plots display gene set scores; the color displays aging type. Adjusted Bonferroni p-values from a Wilcox test are displayed under significant differences at each age. (g) Bar plot illustrating gene expression alterations, identified by aggregating the log2 fold changes of differentially expressed genes unique to each cell state across age. The top 10 most discrepant enriched and depleted cell states between Chronological and Combined aging are shown, bars are also colored by aging type.

**Extended Data Fig. 3.**
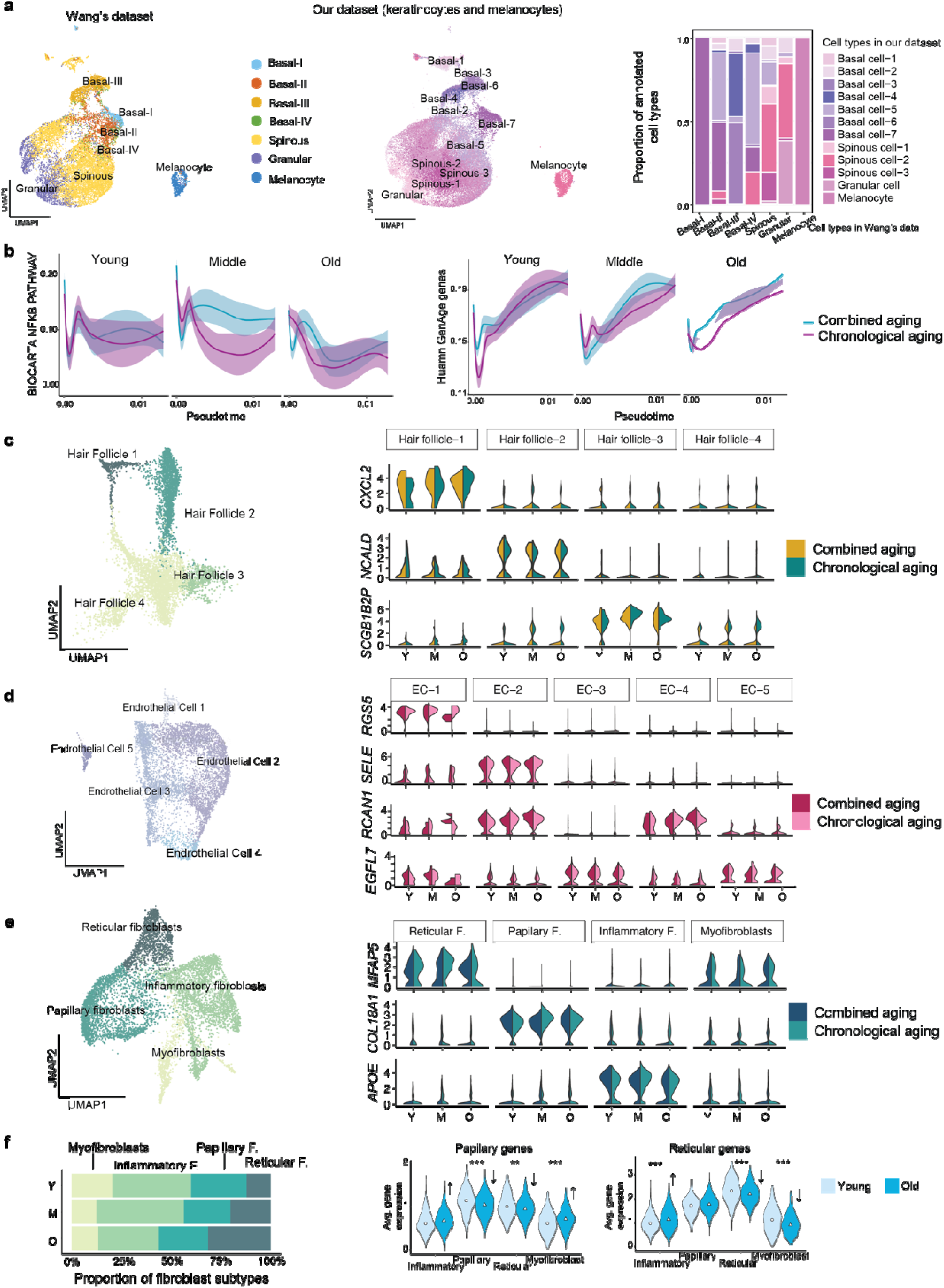
Age-associated transcriptional heterogeneity of human skin cell subtypes. (a) Cross-dataset validation of keratinocyte subtype annotation. Left: UMAP visualization of keratinocytes and melanocytes from Wang et al. Middle: UMAP visualization of keratinocytes and melanocytes from our dataset. Right: Proportional cell type correspondence between Wang’s dataset and ours, confirming high concordance across basal, spinous, and granular hierarchies and validating the robustness of our annotation. (b) Gene expression profiles of NFKB and genAge genes across three age groups and two aging types (combined aging and chronological aging). (c) Analysis of hair follicle subtypes, including subtype categorization and gene expression profiles of marker genes across different samples. (d) Analysis of endothelial subtypes, detailing subtype categorization and gene expression profiles of marker genes across different samples. (e) Analysis of fibroblast subtypes, detailing subtype categorization and gene expression profiles of marker genes across different samples. Left: UMAP visualization of integrated single-cell RNA-seq data from human dermis, identifying four major fibroblast subpopulations: reticular fibroblasts, papillary fibroblasts, inflammatory fibroblasts, and myofibroblasts. Right: Violin plots showing the distribution of representative aging-associated genes (MMP3, COL18A1, and APOE) across the four fibroblast subtypes, comparing samples classified as combined aging and chronological aging conditions. (f) Age-associated changes in fibroblast matrix gene expression. Age-associated changes in fibroblast matrix gene expression. Left: Proportions of fibroblast subtypes in young versus aged skin. Middle and right: Violin plots showing papillary and reticular matrix gene expression across the four fibroblast subtypes in young versus aged samples. **p < 0.01, ***p < 0.001.

**Extended Data Fig. 4.**
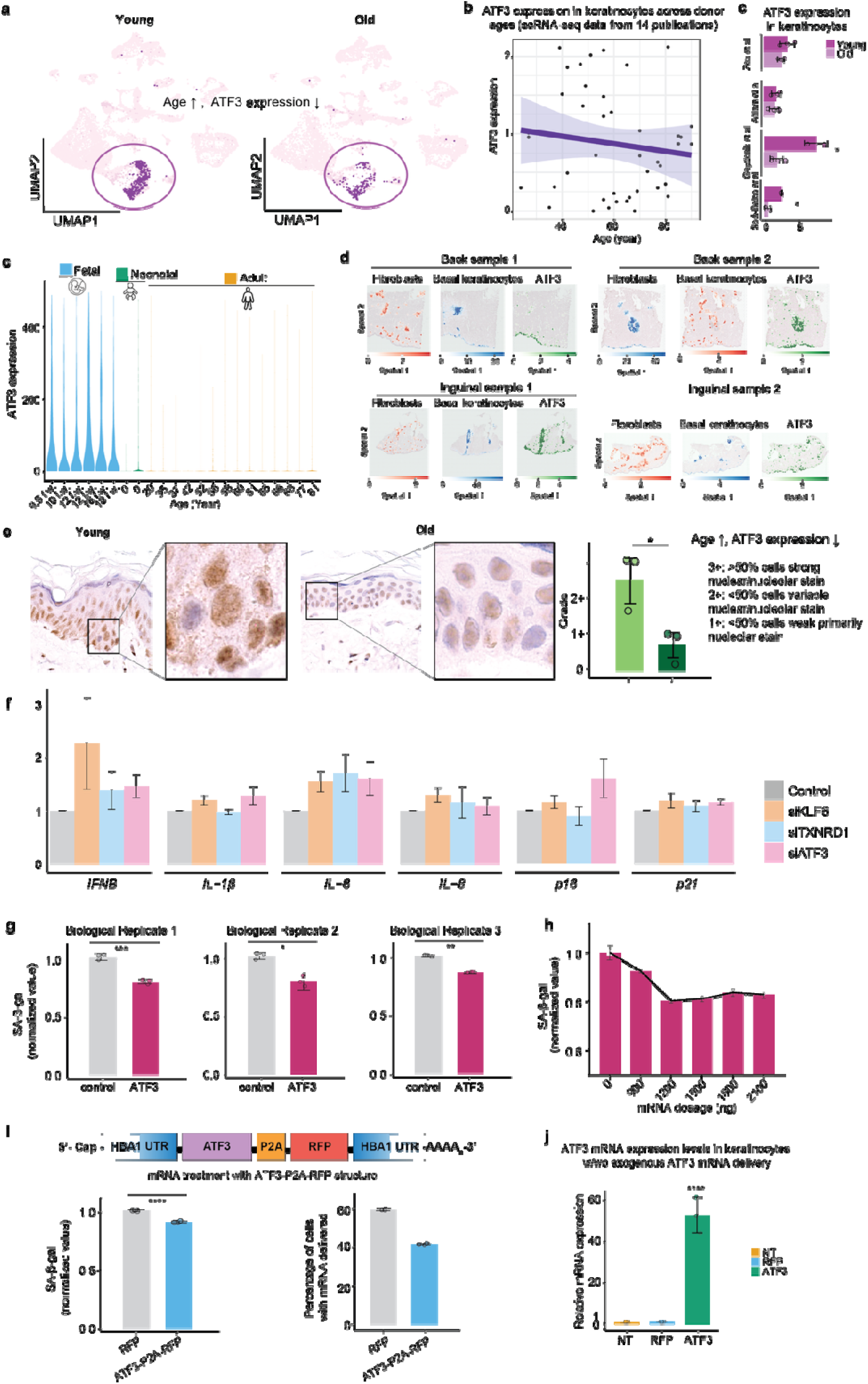
Age-associated decline of ATF3 expression in human skin, functional validation, and therapeutic development. (a) Expression levels of ATF3 in young and old skin samples from our dataset. (b) ATF3 expression in human keratinocytes decreases with donor age in other published datasets. Scatter plot showing ATF3 transcript levels in keratinocytes across donor ages aggregated from 14 publicly available human skin single-cell RNA-sequencing datasets. Each point represents an individual donor sample. The linear regression line (purple) with 95% confidence interval (shaded) reveals a modest but consistent age-associated decline in ATF3 expression (slope < 0). (c) ATF3 expression across epidermal layers at different developmental and age stages. Violin plots showing ATF3 expression in keratinocytes across fetal, neonatal, and adult skin with a progressive decline from fetal to adult stages. (d) Spatial transcriptomics maps of basal keratinocytes (blue), fibroblasts (red), and ATF3 expression (green) in four representative UV-protected body samples derived from the paper Ganier et al., 2024, PNAS. Samples include two from the back and two from the inguinal region. Each set of panels shows fibroblast distributions (left), basal keratinocyte distributions (middle), and ATF3 expression levels (right) overlaid on H&E images. Expression values were scaled to the 99th percentile for visualization. Cell populations annotated in scRNAseq data were computationally predicted on global ST sections using cell2location^33^. Predicted cell abundances shown by color gradients per spot in tissue architecture (H&E) images (representative samples are shown). (e) Immunofluorescence staining of ATF3 in human skin. Scale bar, 200 mm; n = 3 for each group. (f) Quantitative PCR (qPCR) analysis of Senescence-associated Secretory Phenotype (SASP) and senescence-associated β-galactosidase (SA-β-gal) staining of human epidermal keratinocytes upon siRNA-mediated gene knockdown. Data are presented as mean ± SEM. n = 3 per group. **p < 0.01. (g) Comparison of senescence levels in human epidermal keratinocytes treated with mCherry Control and ATF3. The experiments were conducted in triplicate. Senescence-associated β-galactosidase (SA-β-gal) staining using C12FDG revealed significantly lower senescence levels in ATF3-treated cells compared to mCherry-treated controls. Data are presented as mean ± SD, with statistical significance determined by t-test. (h) Dose-response curve of ATF3 mRNA on senescence levels in human epidermal keratinocytes. Five concentrations were tested, and SA-β-gal activity was measured. Data are presented as mean ± 95% confidence interval. (i) mRNA treatment with the P2A structure of ATF3 mRNA. Senescence levels and mCherry positivity were assessed in human epidermal keratinocytes treated with mCherry controls and ATF3 P2A samples. Senescence levels were evaluated using SA-β-gal staining with C12FDG. Data are presented as mean ± SD and statistical significance was determined by t-test. ****p < 0.0001. (j) qPCR analysis of ATF3 mRNA expression levels in keratinocyte with and without exogenous ATF3 mRNA delivery. Data are presented as mean ± SEM. n = 3 per group. ****p < 0.0001.

**Extended Data Fig. 5.**
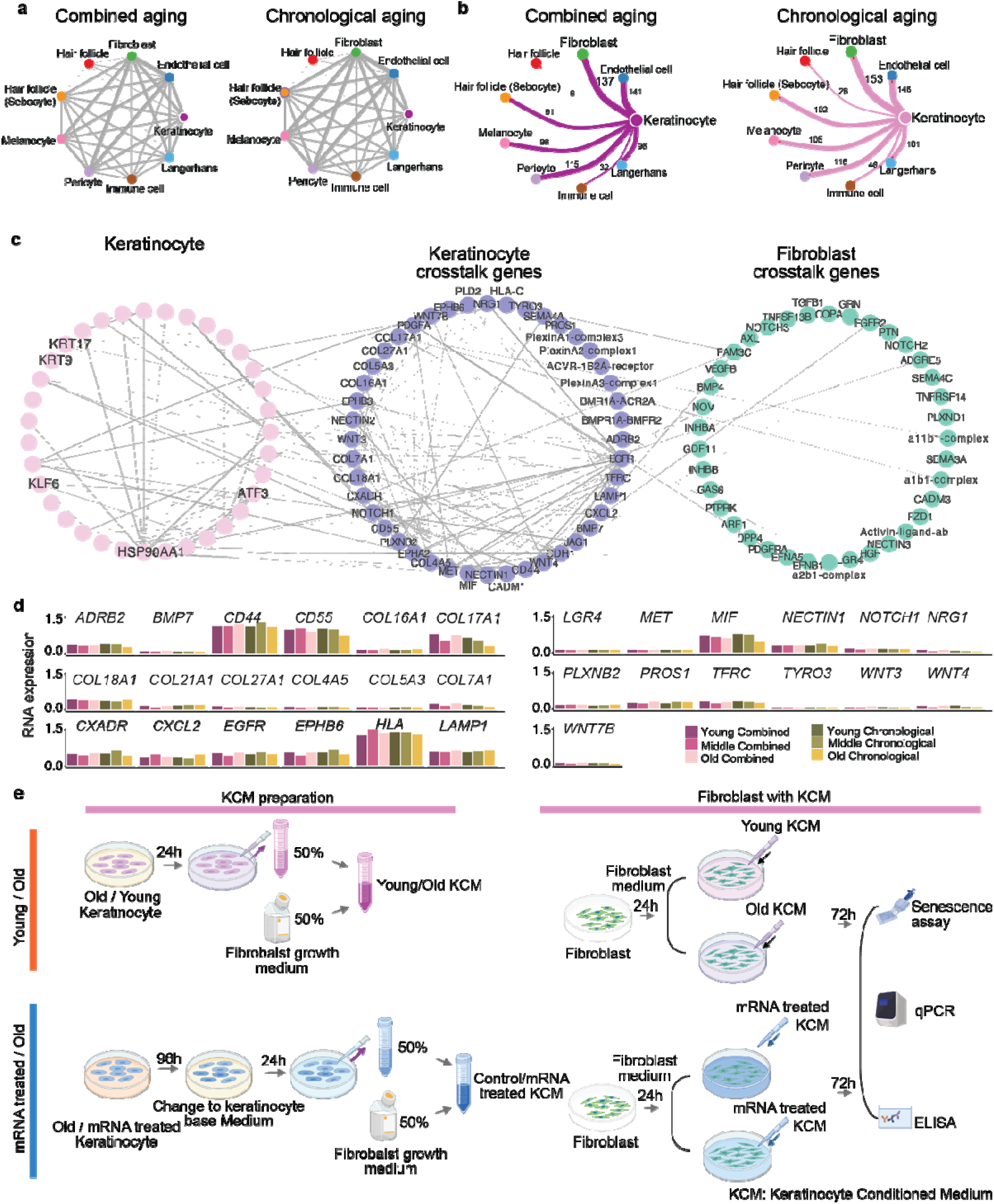
ATF3 mRNA treatment restores fibroblast function through keratinocyte-fibroblast communication. (a) The circo plot illustrates potential cell interactions predicted by CellphoneDB among nine major cell types in both combined aging and chronological aging. These cell types include keratinocytes, fibroblasts, endothelial cells, hair follicles, sebocytes, melanocyte, pericytes, immune cells, and Langerhans cells. The size of each node represents the number ofinteractions, while the width of each edge indicates the number of significant ligand-receptor pairs between the two cell types. (b) The gene-gene network displays potential ligand-receptor pairs between keratinocytes and fibroblasts. (c) The network of gene-gene interactions among down-regulated genes and secreted genes in keratinocytes, as well as the predicted gene-gene connections between keratinocytes and fibroblasts from CellphoneDB. (d) The gene expression profiles of secreted proteins in keratinocytes, involved in the predicted significant ligand-receptor pairs between keratinocytes and fibroblasts, are shown in different samples. (e) Workflow of the Communication Experiment. Keratinocytes were cultured in a basal medium for 24 hours to produce a conditioned medium. This conditioned medium was then mixed 1:1 with fibroblast growth medium and used to culture human fibroblast cells for 72 hours, with fresh conditioned medium replaced every 24 hours. Young KCM, old KCM, control mRNA-treated KCM, and ATF3 mRNA-treated KCM were prepared and utilized for fibroblast culture. After 72 hours, measurements were taken for β-galactosidase activity, collagen and elastin mRNA levels by qPCR, and collagen levels by ELISA.

**Extended Data Fig. 6.**
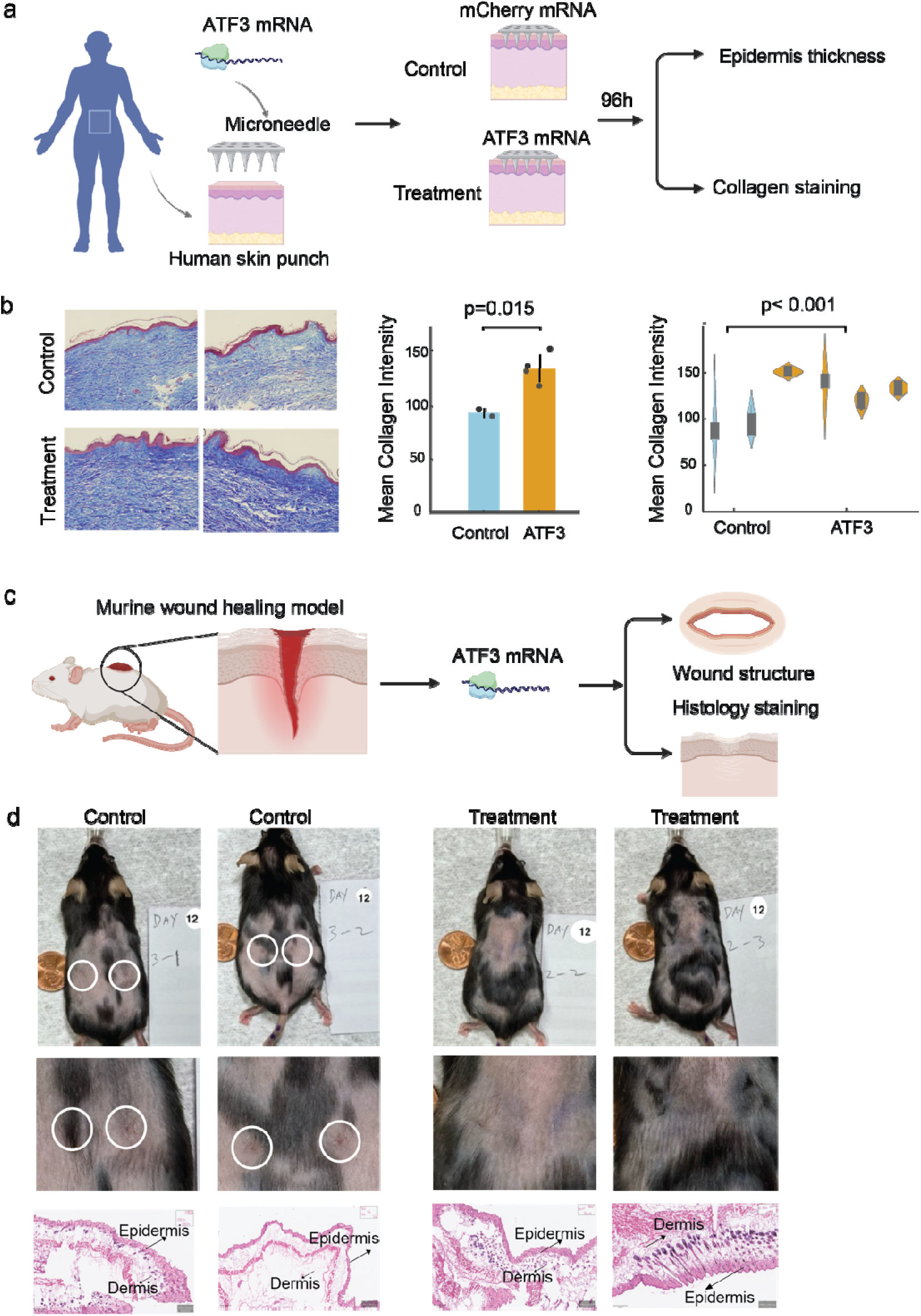
ATF3 mRNA rejuvenation validation in *ex vivo* human skin and *in vivo* murine model. (a) Schematic illustration of the experimental setup showing microneedle-assisted delivery of lipid nanoparticle–encapsulated ATF3 mRNA or control RFP mRNA into 61-year-old human skin punches cultured *ex vivo* for 96 h. (b) Representative histological and immunofluorescent images showing increased epidermal thickness and collagen staining following ATF3 mRNA treatment compared with control mRNA. Scale bars, 100 µm. (c) Quantification of mean collagen intensity and epidermal thickness (mean ± s.e.m.) in *ex vivo* human skin, showing significant enhancement upon ATF3 mRNA treatment (p = 0.015). Statistical significance was assessed using unpaired two-tailed t-tests. (d) Representative images of the murine full-thickness wound healing model demonstrating improved wound structure and collagen remodeling after ATF3 mRNA delivery.

